# Comparative Analysis of Gene Importance in *Escherichia coli* Across Growth Conditions

**DOI:** 10.1101/2025.10.02.679994

**Authors:** Antoine Champie, Simon Jeanneau, Amélie De Grandmaison, Mathias Martin Silva, Jean-Philippe Coté, Pierre-Étienne Jacques, Sébastien Rodrigue

**Author notes:** **Correspondence** To whom correspondence should be addressed: Sébastien Rodrigue, Université de Sherbrooke, Sherbrooke, 2500 boulevard de l’Université, Sherbrooke, QC J1K 2R1, Pierre-Étienne Jacques, Université de Sherbrooke, Sherbrooke, 2500 boulevard de l’Université, Sherbrooke, QC J1K 2R1. These authors contributed equally to this work (Antoine Champie, Simon Jeanneau, Amélie De Grandmaison).

## Abstract

As the ability to synthesize complete genomes becomes increasingly accessible, the question of what should compose those sequences is becoming more prevalent. However, identifying genes essential for the survival of an organism is challenging, as gene essentiality is a nuanced concept that heavily depends on context. In this study, we identified growth medium-specific important genes by performing transposon mutagenesis in *Escherichia coli* BW25113 and sequencing mutant populations at multiple time points in three growth media. Our analysis revealed a core set of 412 important genes shared across all conditions, along with distinct medium-specific gene sets of varying sizes. By analyzing temporal variations in read counts for each gene, we identified additional sets of genes whose inactivation causes a lighter impact on fitness. We used this dataset to define medium-specific gene modules required to sustain robust growth under each condition. Our study underscores the context-dependent nature of gene essentiality and marks a step toward refining the concept from a universal list to a more nuanced, condition-specific framework, which will be invaluable for future genome design efforts.

**Graphical abstract:** 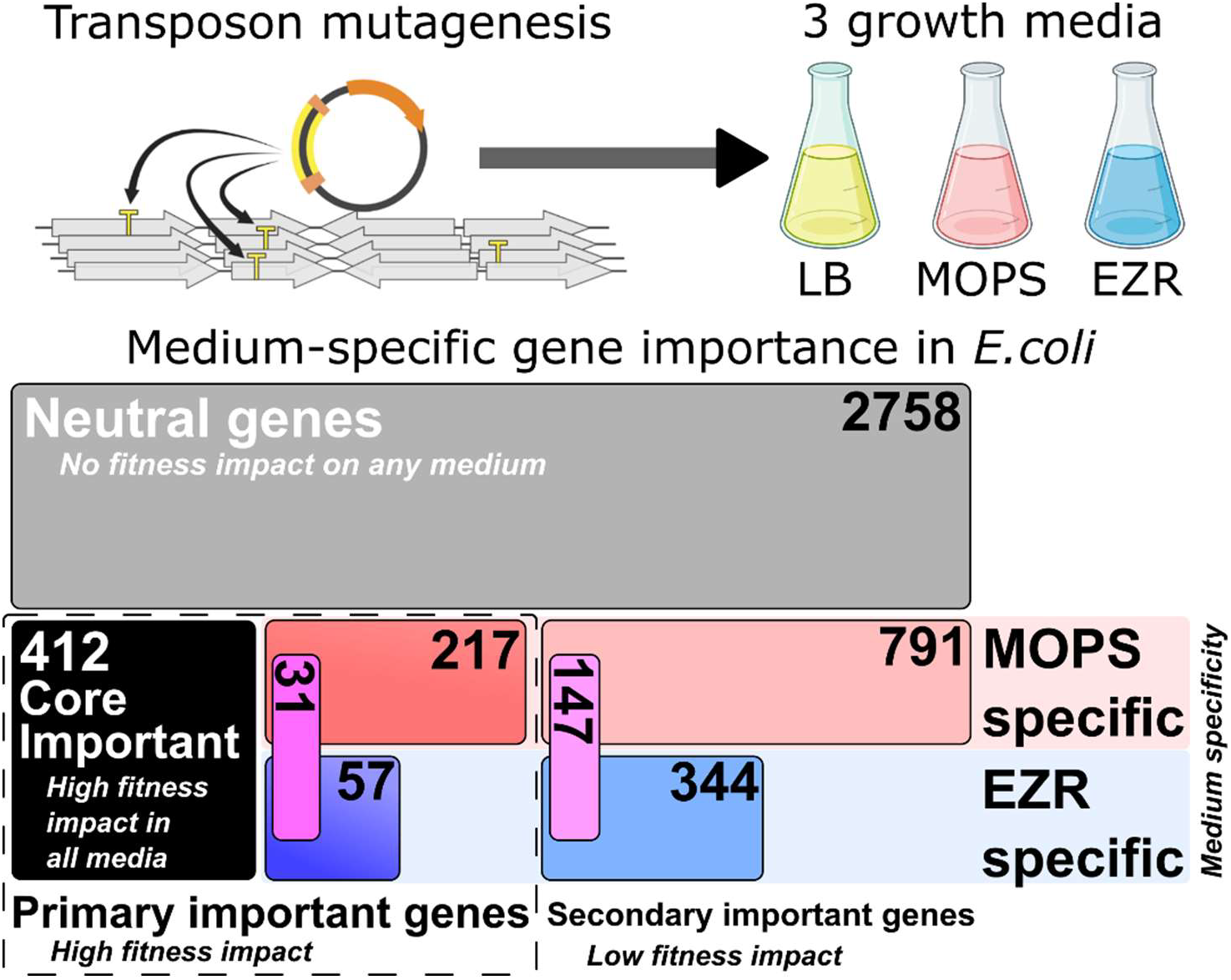

## Introduction

As methods for genome synthesis and assembly improve^1–3^, it is becoming more important to consider what sequences should be included in a synthetic genome^4^. While general knowledge of biological mechanisms steadily increases over time, our comprehension is still far from exhaustive. For example, despite decades of research, the role of approximately 35 % of *Escherichia coli* genes remains elusive^5,6^, and their importance for robust growth is undetermined in many cases. This knowledge gap is particularly problematic for genome minimization, which involves removing non-essential elements from a genome while preserving its viability and functionality, since a single error in genome composition can compromise the viability of the entire organism. Even a decade after the synthesis of the first minimal genome^7,8^, which was based on the naturally small chromosome of *Mycoplasma mycoides*, research is still ongoing to fully elucidate the function of all its genes^9^. Replicating this achievement in more complex organisms remains a difficult task.

While a relatively small group of genes encoding key functions, such as DNA replication and gene expression, is likely to be strictly essential under any circumstances, the importance of many genes depends on environmental conditions. Metabolic genes, for example, are known to display differential importance depending on the composition of the growth medium. This variability underscores the importance of understanding how environmental factors influence gene essentiality, as such insights will be crucial for tailoring novel genomes for specific media and conditions.

Extensive data is already available to evaluate gene essentiality in well-studied microorganisms. In *E. coli*, for instance, different methods such as comparative genomics^10^, genome-scale models (GEMs)^11^, targeted gene deletions^12^, transposon insertion sequencing (TIS)^13–17^ as well as CRISPR interference (CRISPRi)^18^ have been employed, each providing a unique perspective on the question. However, significant challenges exist in integrating these datasets to accurately quantify the conditional importance of a given gene in this bacterium. One of the primary barriers to accurately comparing different datasets is the wide variation in growth conditions used to selectively grow gene-inactivated mutants. These differences encompass various factors, including medium composition, clonal versus bulk population cultures, temperature, aeration level, and the growth phases included in the experiments. Another barrier is the various strategies and thresholds employed to assign an essentiality status to a gene^19–23^. Finally, the specific strains used to generate gene essentiality datasets can also vary, which may also cause discrepancies due to differences in genetic backgrounds. Ultimately, even if the data available on gene essentiality is particularly extensive in *E. coli*, pinpointing the causes behind observed variations between studies remains a complex task.

In an attempt to reliably measure and compare the impact of gene inactivation under different growth conditions, we performed transposon mutagenesis in three different media using *E. coli* BW25113, the parental strain of the Keio collection of single-gene deletion mutants^12^. The “MOPS minimal glucose” and “EZ-Rich glucose” defined media, respectively abbreviated MOPS and EZR, were selected as they were specifically developed for *Enterobacteriaceae* growth^24^. MOPS minimal medium is a growth medium buffered using morpholino propane sulfonate (MOPS) and Tris in which only glucose, ammonium, and base levels of metallic ions are provided. EZR medium is a MOPS medium in which all nucleic acids, amino acids, and some vitamins are also provided in defined concentrations. To search for genes required for metabolism beyond glycolysis, the primary driver of the Krebs cycle in the first two media, we also included LB medium. LB is a commonly used, chemically undefined medium in which metabolism relies primarily on proteolysis^25^. In this context, a variety of genes beyond those involved in glycolysis are required to catabolize amino acids into precursors for central metabolism. In addition, several gene essentiality datasets already exist in LB, notably those derived from the Keio collection^12^ and recent TIS experiments^14^, thus facilitating comparison with our results.

## Results

### Transposon mutagenesis in LB, MOPS, and EZR media

We applied the recently developed high-throughput transposon mutagenesis (HTTM)^15^ method to generate 60 independent mutagenesis replicates in *E. coli* BW25113. The initial mutated populations (P0) were then divided into three groups of 20 replicates for growth in LB, MOPS, or EZR for five consecutive daily passages, referred to as P1 to P5 (**Fig. 1A**). As observed in previous genome streamlining endeavors, a genome composed solely of essential genes is likely to exhibit significant fitness defects and limited applicability in biotechnology^4,26^. This highlights the need to conserve additional genes that contribute to maintaining a robust organism, commonly referred as fitness genes or quasi-essential genes^26,27^. To ensure detection of all genes important for bacterial life (both strictly essential genes and major fitness contributors), the mutants were left to compete against the rest of the population, resulting in progressive depletion of the mutants with suboptimal fitness. The populations were also intentionally allowed to grow in stationary phase to eliminate mutants that were not able to transition efficiently between different growth phases. Because this approach identifies all genes whose inactivation affects the relative fitness of the strain across different growth phases due to various factors, we categorized them under the broader term ‘important genes’ rather than using the traditional ‘essential genes’ terminology.

**Figure 1.**
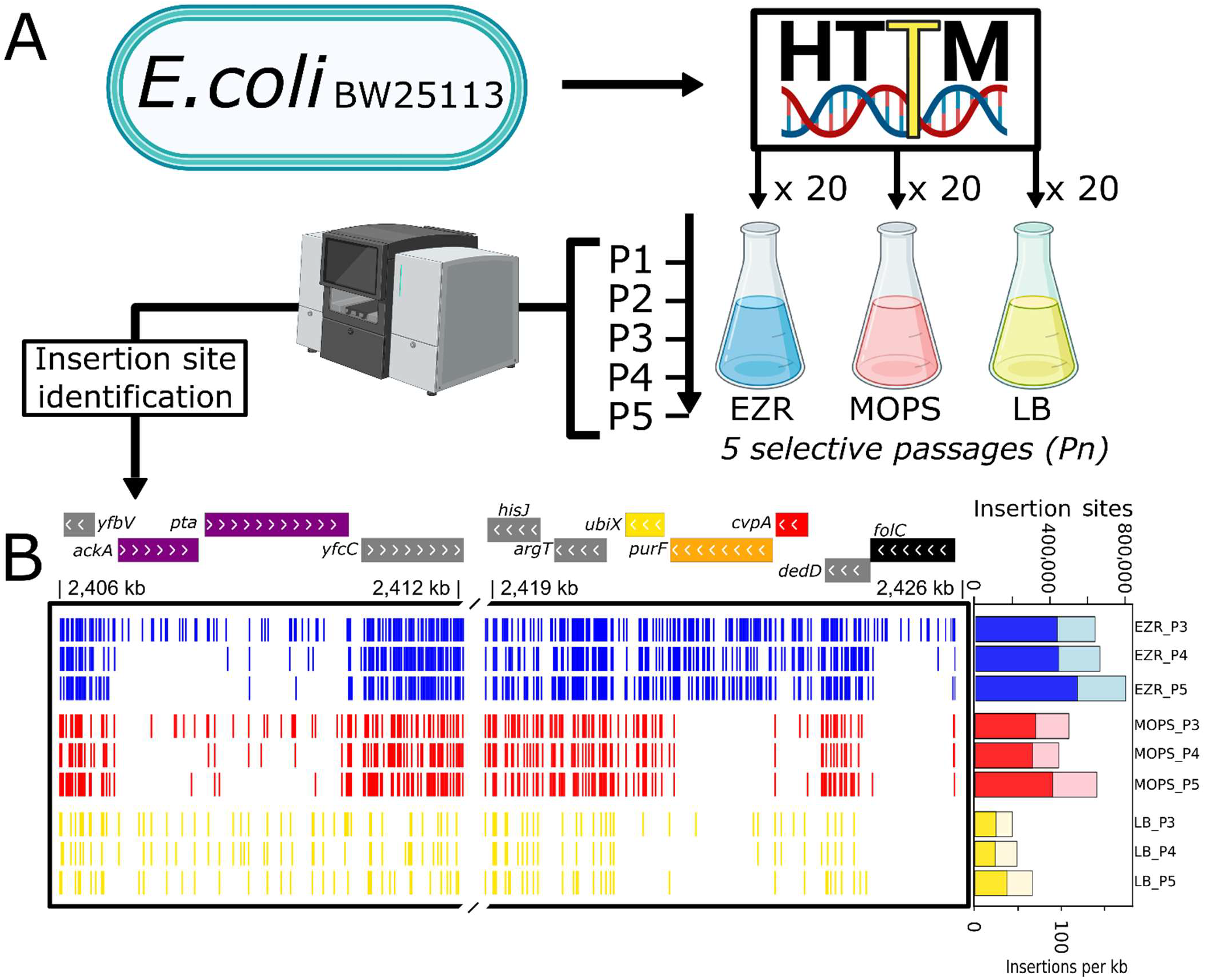
Overview of the experimental strategy used in this study. **A)** Schematic description of the transposon mutagenesis experimental procedure. A total of five culture passages were performed for each medium (P1 to P5). **B)** Representative genomic loci displaying mapped transposon insertion sites across passages three to five in each tested culture medium. Genes with different transposon insertion densities such as *yfcC* (densely riddled by transposon in all three media), *pta* (showing a gradual reduction of insertions over passages), and *folC* (considered essential in the Keio collection and devoid of transposons in all three conditions at P5) can be regrouped according to the medium or media in which they have been determined important (gray = non important, black = important in all media, yellow = important in LB only, orange = important in LB and MOPS). The total number of insertions detected per passage among all pooled replicates is displayed on the right. Paler coloration indicates total insertions found, whereas solid color indicates genic insertions.

To observe the gradual depletion of transposons in moderately important genes over time, as recently observed in *Acinetobacter baylyi*^28^, the remaining cells were harvested after each passage (P0–P5), frozen, and their DNA extracted. Samples were then prepared for next-generation sequencing and the transposon insertion sites were identified as previously described^15^. All 20 replicates of the final passages and twelve replicates of each intermediate passage (P0-P4) were sequenced for each medium, totaling 180 sequenced samples.

Early tests revealed an overwhelming predominance (> 95 %) of reads originating from the mutagenesis plasmid pFG051 in early passages (P0-P2), resulting in very low genomic insertion counts (see **Table S1**). To mitigate this phenomenon, we implemented an additional restriction enzyme-based depletion step to the previously described library preparation process^15^ (see Materials and Methods for additional details) which reduced plasmid-associated reads to an acceptable level (**Table S1**).

Reads from all replicates for each condition were then pooled and used to identify transposon genomic insertion sites. This yielded a considerable number of insertions (at P5; EZR = 837,713; MOPS = 650,513; and LB = 306,072) (**Fig. 1B, Fig. S1 and Table S1**). Considering the genome size of *E. coli* BW25113, this corresponds to an average of one insertion every 5.5 bp, 7.7 bp, and 20.0 bp in EZR, MOPS, and LB medium, respectively. The location of those insertions can be visualized through the provided UCSC genome browser public track hub (https://genome.ucsc.edu/cgi-bin/hgHubConnect?hgHub_do_redirect=on&hgHubConnect.remakeTrackHub=on&hgHub_do_firstDb=1&hubUrl=https://g-f2b62d.6d81c.5898.data.globus.org/Champie_2025/Champie_2025.hub.txt)

### Definition of a gene-importance metric

Many different methods have been described for analysing transposon mutagenesis data and assigning an essentiality status to genes ^20–22,29,30^. These approaches usually rely on the ability to discriminate between gene populations with high vs low numbers of transposon insertions using various statistical tools. Different metrics can be used to quantify the number of genic insertions, simple ones such as reads per base and insertions per base are convenient but not particularly robust, as the former is particularly sensitive to sparse insertions supported by abnormally high number of reads, while the latter is more affected by background noise insertions characterized by very low read counts.

Given the high transposon insertion density and sequencing depth of our datasets, we reasoned that alternative strategies leveraging both read counts and insertion site numbers could be explored to give a more nuanced estimation of gene importance, while mitigating the aforementioned pitfalls. Therefore, we developed a new method that leverages both read and insertion site numbers to quantify the proportion of each gene harbouring only background levels or no insertions, which is then used as a proxy to estimate the importance of each gene for cell fitness. Briefly, all genes are scanned using overlapping sliding windows (bins) whose parameters are adjusted for each sample. The bin size is determined using the average distance between transposon insertion sites, and the threshold is calculated based on the median number of reads per insertion. Each bin is then evaluated and classified as a HIT (*Highly Inserted segmenT*) or a MISS (*Minimally InSerted Segment*) depending on whether the total number of reads is above or below the defined threshold. (**Fig. 2A, Fig. S2**). A gene-importance score (*i*Score) is then calculated based on the proportion of *MISS* bins, where a score of 100 % would represent an important gene that does not tolerate any significant level of transposon insertions anywhere on its sequence. The distribution of *i*Score per gene is initially almost uniform for all experimental conditions, and splits into a clear bimodal distribution as the number of passages increases (**Fig. S3**). The left mode of this final distribution consists of genes of low importance and can be modelled using an exponentially decreasing distribution. For each medium, genes were defined as “important” if they had less than a 0.01% probability of belonging to this low-importance population, according to the fitted distribution (**Fig. S4**). The complete *i*Score dataset for the last passage in all media is available in **Table S2**.

**Figure 2.**
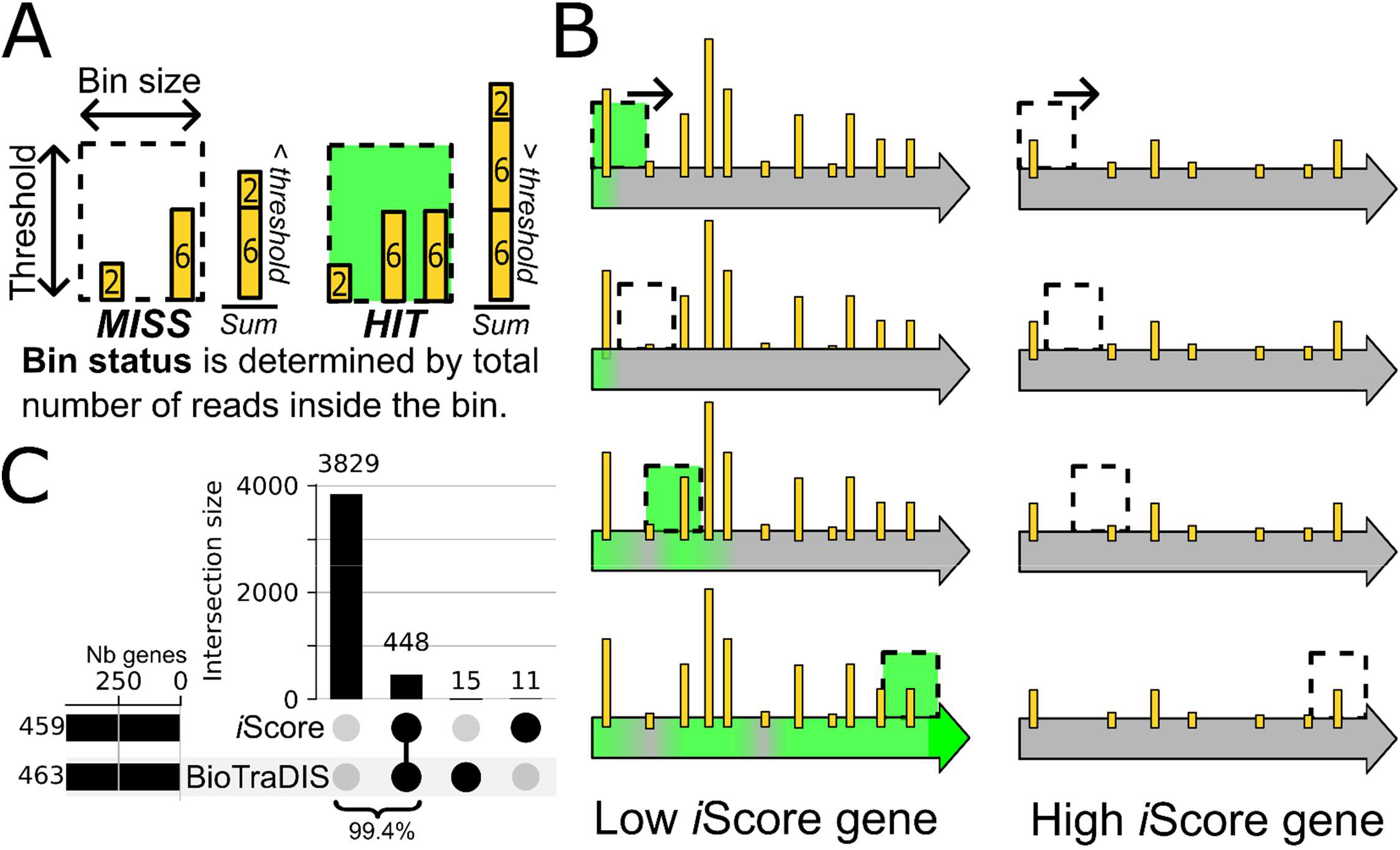
Bin processing of transposon insertion sites. **A)** Details on data-driven determination of bin parameters. A bin is labeled as “*HIT*” if the number of reads exceeds the threshold and otherwise labeled as “*MISS*”. The threshold and bin size for each condition are available in **Figure S3.** Yellow bars represent single base pair-resolution transposon insertion sites, and the height of the bars represents the number of mapped reads at a given position. **B)** Graphical representation of the gene scanning procedure and *i*Score attribution. **C)** Comparison of the number of essential genes (black circles) in EZR after Passage 5 with data processed using either *i*Score or the Bio-TraDIS^21^ methodology.

The comparison of this new approach to the well-established Bio-TraDIS tool^21^ has shown a very high agreement rate for MOPS and EZR media (98.3 % and 99.4 %, respectively), while the concordance in LB medium was lower (91.3 %) but still very high (**Fig. 2C, Fig. S5)**. Those genes, hereafter referred to as primary important genes, were grouped according to their status in each medium, with genes important in all three media forming the core important genes.

To validate our classification, we compared our primary important gene lists to metabolic modelling predictions. For EZR (**Fig. 3A**) and MOPS (**Fig. 3B**) media, we used the *i*ML1515 metabolic model, simulating each medium independently for comparison^31^. However, since LB is a complex and undefined medium, simulating its composition is prone to errors. The results for LB (**Fig. 3C**) were therefore compared with existing high-quality essentiality datasets generated using this medium^12,14^. We observed a strong agreement in all three cases, with a general trend of detecting most previously known genes, as well as additional genes. This trend was expected, considering the high sensitivity of our approach to reduction in fitness. For example, the genes composing the *nuo* operon, all of which were previously deleted in the Keio collection and successfully cultivated in LB^12^, are found to be of primary importance only in LB medium in our dataset (**Fig. 3D**). Competition assays in the selective conditions used for our HTTM experiment confirmed that the Δ*nuoF* mutant is quickly outcompeted by the parental strain in LB medium only (**Fig. 3E**).

**Figure 3.**
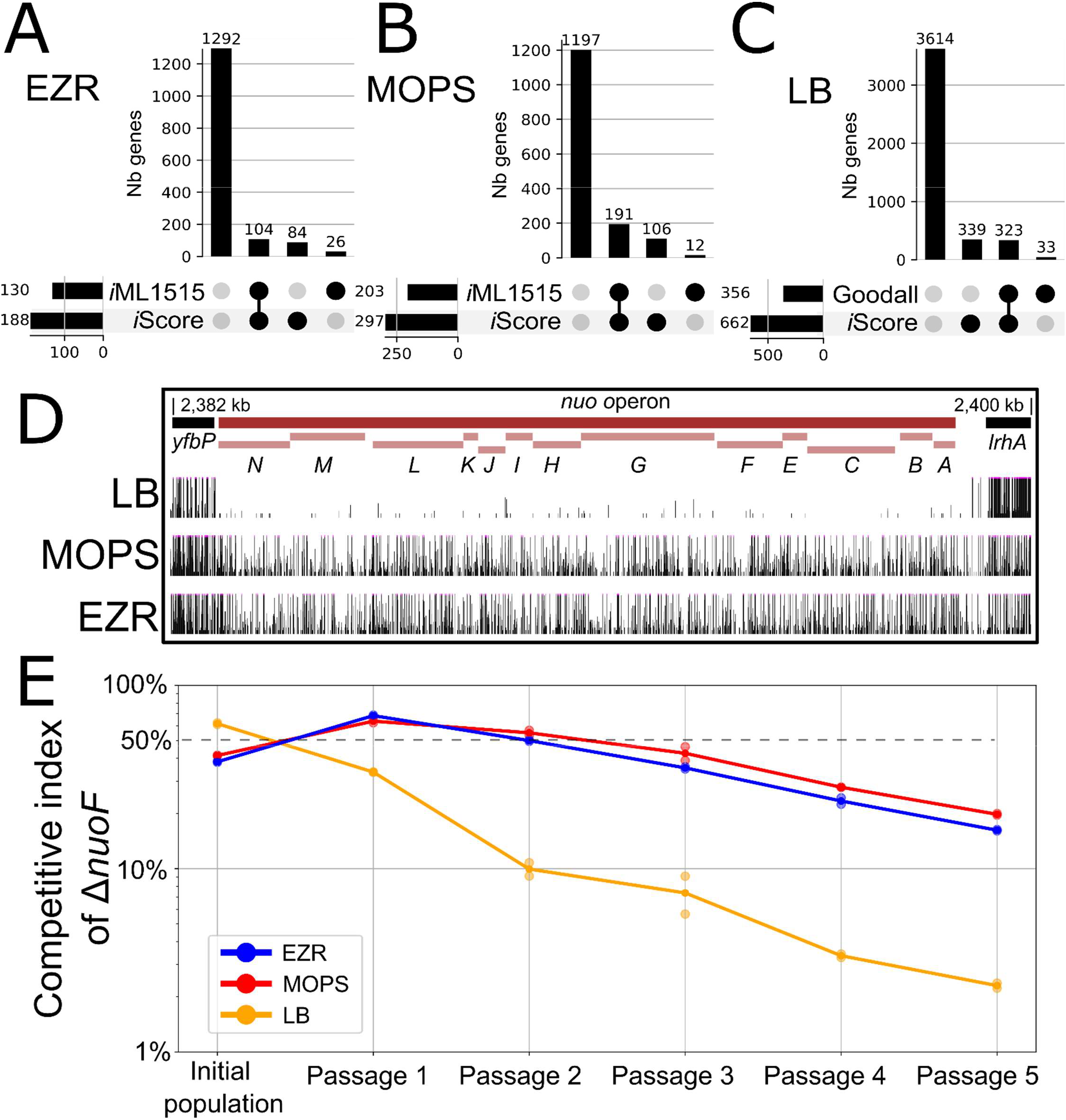
Comparison of essentiality calls with existing datasets. **A-B)** Comparison between genes predicted to have a significant impact on fitness by the *i*ML1515 model and our data in EZR (A) and MOPS (B). Black circles indicate primary important or essential genes. **C)** Comparison of LB primary important gene calls determined in our study (*i*Score) with previously published LB essential genes (E-Genes from Goodall *et al*^14^). **D)** Representative locus showing an operon (*nuo*) with differential importance between tested culture media. **E**) Competitive assay between *E. coli* BW25113 and BW25113Δ*nuoF* performed in all three media over five passages.

### Characterization of core important genes

Most of the primary important genes are common in all three media (**Fig. 4AB**). As expected, these core important genes are generally responsible for the basal functions of the cell required regardless of medium composition, such as genetic information processing or glycan production, based on the KEGG database classification^32^ (**Figs. 4C, 4D**). Some metabolic genes are also found in the core, for example genes associated with the biosynthesis of lipids, which are not provided in any of the three tested media (**Table S4**). Interestingly, only two genes were classified as important in the three media while belonging to the “Poorly characterized - Function unknown” category in the KEGG database. Comparison with the Ecocyc database, a more comprehensive and regularly updated source for gene characterization^5^, revealed a more accurate count of 15 uncharacterized genes and 32 partially characterized genes (**Table S2**), highlighting targets that warrant further investigation. With the core set of important genes defined, we next focused our investigation on the subset of genes specifically required in the defined MOPS and EZR media.

**Figure 4.**
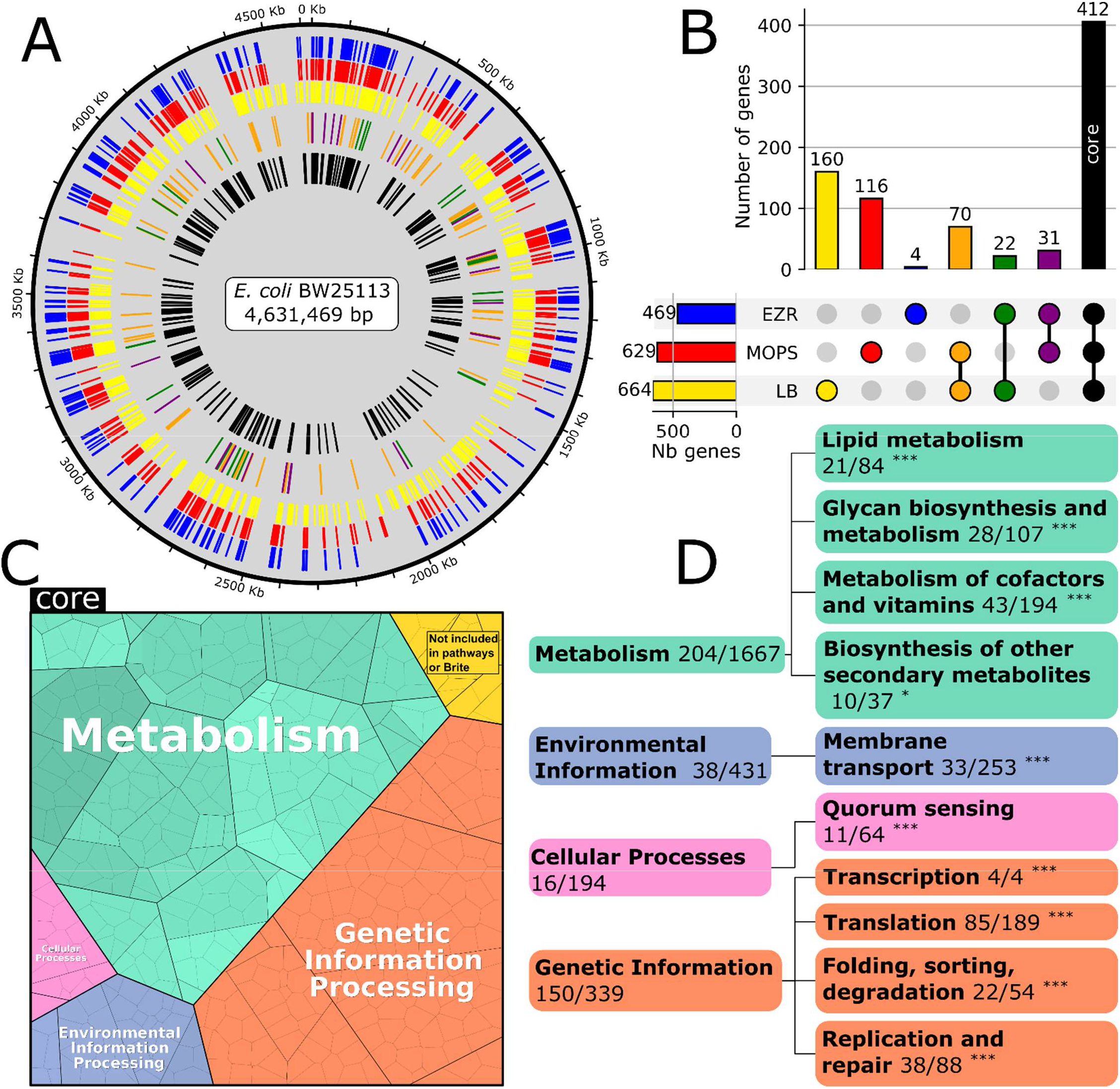
Description of the core important genome. **A)** Repartition of the important genes on the *E. coli* BW25113 chromosome. Each color in the outer rings indicates importance in a condition (EZR = blue, MOPS = red, LB = yellow), bars in the middle ring indicate importance in 2 media (color matching), and black bars in the inner ring depict genes important in all 3 conditions (core). Colors are constant across the article. **B)** Upset chart representing the size of the different subsets of important genes. **C)** Repartition of the functions of the core important genes into the main functional categories of the KEGG database, represented using the Proteomaps tool^33^. An interactive version is available at (http://bionic-vis.biologie.uni-greifswald.de/result.php?jobID=17097408404332&version=UserSpec). **D)** Decomposition of the main functional categories of core important genes. The significance of ontology term enrichment compared to random expectation was tested using a hypergeometric distribution; p-values are indicated as follows: * < 0.05, ** < 0.01, *** < 0.001.

### Detection of important regions in genes

To refine the detection of important genes, we sought to identify genes that possess an important region but can tolerate insertions in the rest of their sequence. Such genes could have a relatively low *i*Score but would still result in major fitness loss upon removal. We identified these genes by detecting stretches of contiguous MISS bins within genes exhibiting intermediate iScores (15 < iScore < 90), in combination with a clear split point where most bins are labelled *HIT* on one side and *MISS* on the other. Cases such as *ftsN* or *mqsA* (**Fig. 5AB**) are straightforward to interpret, with the important protein section matching known *ftsA* interaction domains^34^ or a toxin (MqsR) interaction domain^35^, meaning that the 3’ of these genes are likely dispensable for the genes’ functions in these growth conditions. Other genes, such as *gpt*, which encodes a xanthine-guanine phosphoribosyltransferase, appear to have an important section only in EZR (Fig. 5C). We speculate that the role of its C-terminal region as a purine salvage enzyme is vital for competitiveness only in EZR. Using an automated approach followed by manual curation, we identified 57 and 55 genes that harbor at least an important region that should be conserved for growth in MOPS and EZR media, respectively. The complete list is available in **Table S5**.

**Figure 5.**
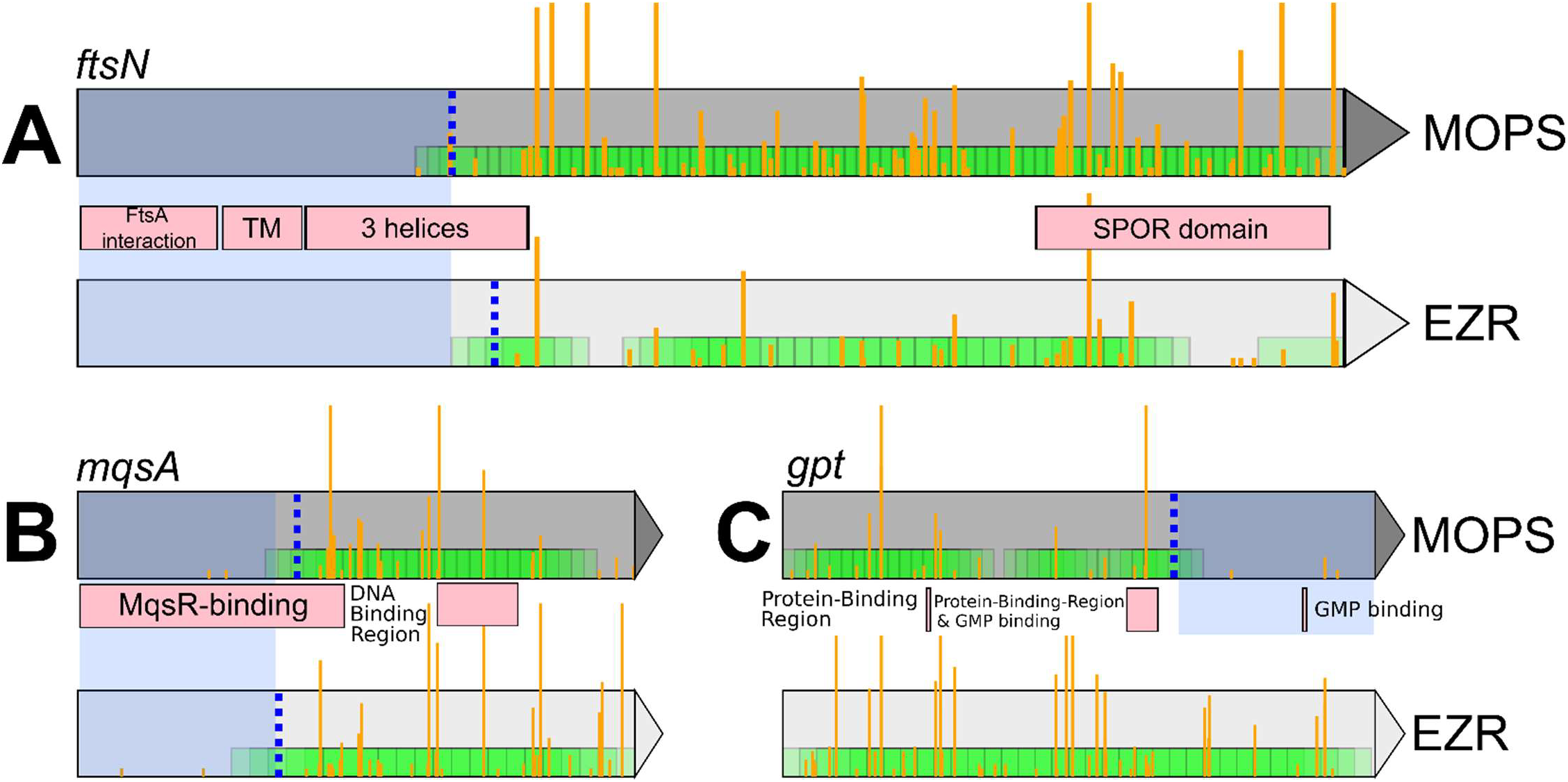
Examples of genes with an important region in MOPS or EZR media. A-C) *ftsN, mqsA*, and *gpt* genes, respectively. Yellow bars indicate transposon insertion sites, with height proportional to the number of mapped reads. The dashed blue line represents the optimal segregation between the *HIT* bin (Green) and the important region (highlighted in blue). Pink boxes represent known domains and features (Uniprot database).

### Identification of secondary important genes using temporal variation

In HTTM experiments, an insertion mutant that carries even a slight fitness burden is expected to be gradually outcompeted by the rest of the population, which is reflected by a continual decline in associated read count^28^. Conversely, an unaffected insertion mutant would have a constant or even increasing read count over the passages. Primary important genes identified using the *i*Score approach reached their lowest read count at P5 or earlier, reflecting a strong impact on cell fitness upon inactivation. We then investigated the remaining non-primary important genes to assess their influence on fitness. Based on hierarchical clustering of the read fluctuation over time in each media, we performed a qualitative segregation of genes into three nuanced groups: secondary important genes that would be likely devoid of insertions after additional passages, precarious genes have an ambiguous status where they may lightly impact the fitness upon inactivation, and neutral genes who can be removed with minimal impact (**Fig. 6, Fig. S6 and Table S6**). Depletion rates vary widely among genes of neutral importance, ranging from nearly null to strongly positive. Those latter atypical cases might reflect an effective fitness gain when inactivated or simply reflect an effect of the mere compositional nature of the data. Secondary important genes were appended to the previously determined list of primary important genes to form medium-specific genes modules containing the complete non-core gene set required for robust survival in either MOPS or EZR (**Fig. 7A**).

**Figure 6.**
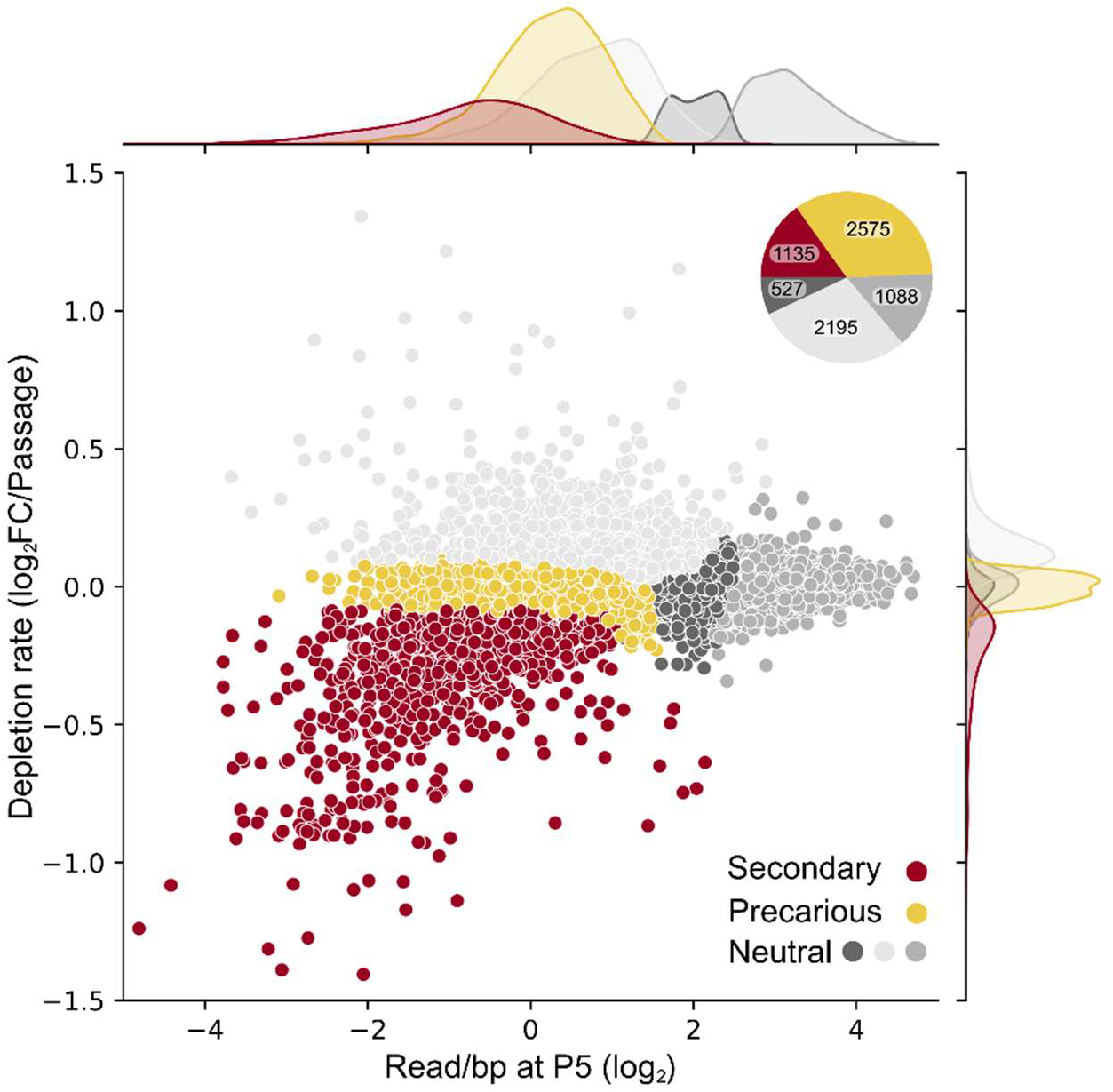
Temporal variation of insertion counts uncovers fitness affecting genes. Depletion rates of normalized read counts across passages in both defined media are shown for all non-primary important genes vs their total read count per base at P5. Values for both EZR and MOPS are plotted as individual dots. Genes are colored according to hierarchical cluster assignments (secondary, precarious, or neutral). Marginal density plots along the axes show the distribution of each metric across gene categories. The inset pie chart displays the absolute gene count per category, both medium combined (see **Fig. S6** for extended view including primary genes).

**Figure 7.**
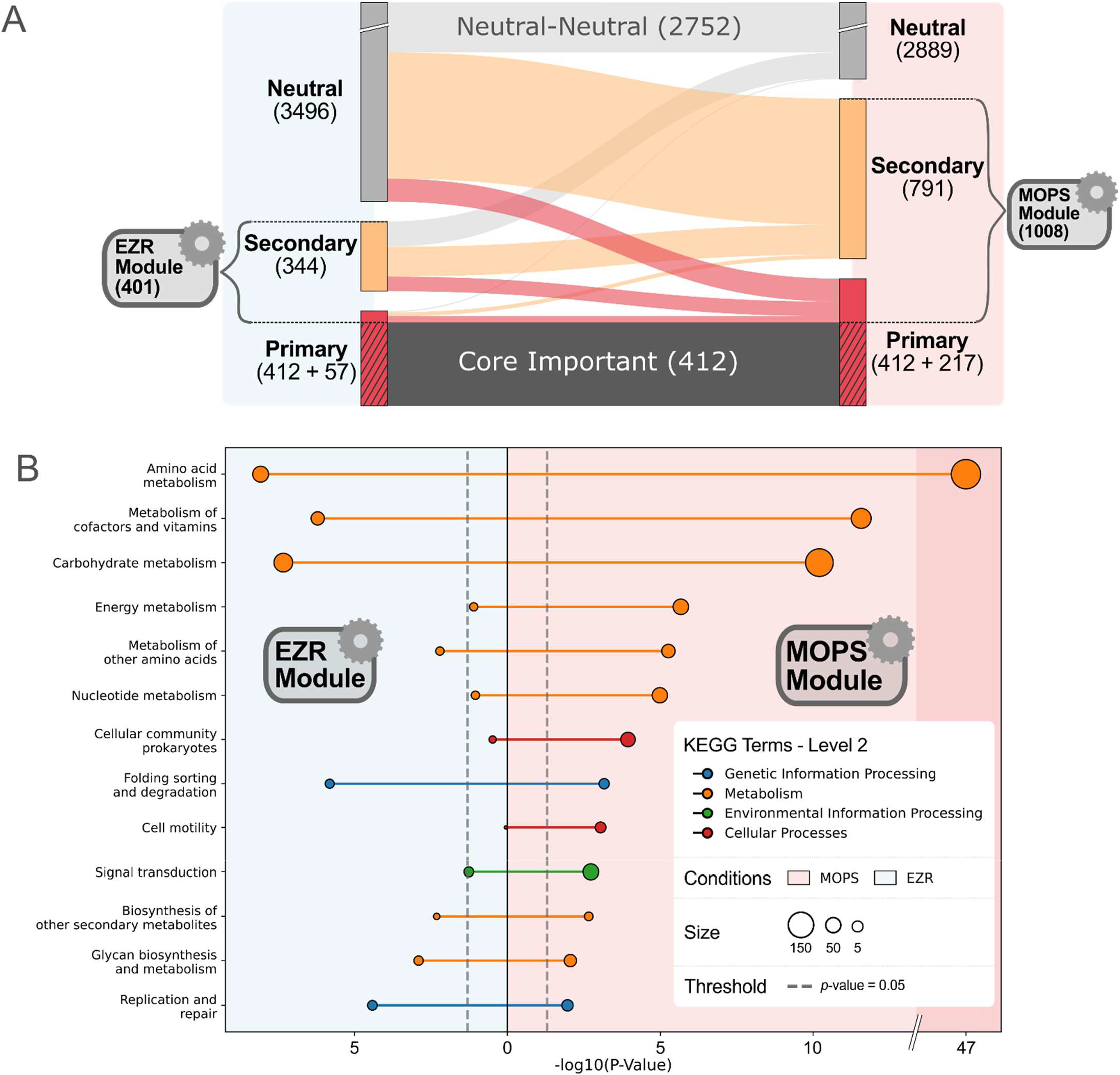
Description of the medium-specific gene modules. **A)** Sankey diagram showing gene status in EZR (left) and MOPS (right) media. Flows indicate changes in gene importance between conditions, with colors (gray, orange, and red) reflecting the importance category in MOPS. Numbers denote the gene counts for each category and module. **B)** Lollipop chart representing Level 3 KEGG ontology enrichment analysis for genes in the EZR and MOPS modules. The -log_10_(p-value) is plotted for each ontology term, with higher values indicating stronger enrichment. Only ontology terms with a p-value below 0.05 are shown.

### MOPS module

In total, 629 genes were classified as primary important for growth in MOPS medium. Among these, 412 are shared across all conditions, the previously defined core important genes, while 217 are specifically important in MOPS. In addition, 791 genes were classified as secondary important in this medium. Together, these two groups define the MOPS module, comprising 1,008 non-core important genes **(Fig. 7A, Table S7)**. Functional analysis of genes important for growth in MOPS revealed significant enrichment in metabolism-related genes, as expected from the composition of this minimal medium (**Fig. 7B**). Notably, pathways responsible for the biosynthesis of all 20 standard amino acids and all four nucleotides exhibited significant enrichment (Bonferroni-corrected p-value < 0.05, **Table S4**). The mapping of the MOPS module gene set onto the KEGG “Biosynthesis of Amino Acids” metabolic pathway confirms that all reactions required to synthesize the 20 amino acids from central metabolic precursors are covered (**Fig. S7**). Notably, some of the reactions in these pathways can be performed by two or more genes, which should be individually less susceptible to inactivation. In each of these cases, one of the redundant genes is identified as a secondary important gene with the single exception of the last step of asparagine synthesis. This final step can be catalyzed by either AnsA or AnsB^36^, two enzymes that appear capable of fulfilling the cell’s asparagine requirement independently, without measurable loss of fitness.

Predictions of the fitness impact of single-gene inactivation were performed using the *i*ML1515 metabolic model under MOPS-glucose medium conditions (**Fig. 3B)**. This comparison revealed a high degree of concordance between the two approaches (∼ 92.1 %). The 12 genes that were predicted as important only by the model (*luxS, pabA, ubiD, ispA, bioH, panB, yrbG, zupT, hemG, hemL, pabB and panC*) are all long and densely inserted genes. Those genes might be functionally complemented by other mutants of the population or simply mis-predicted as important by the model. Among the hits not consolidated by the model, we identified some generic information-processing genes, such as *lysS* (tRNA synthetase). The high *i*Score of this gene in defined media (100 in MOPS, 84.8 in EZR) suggests that its removal would severely impair the strain in these conditions, while having only a moderate impact in LB medium, where the *i*Score is 25.6. The reason why *lysU*, a gene known to perform the same function^37^, does not equally compensate under all media conditions remains unclear.

### EZR Module

The EZR module is composed of 401 non-core genes, among which 57 are primary important and 344 secondary important in this medium (**Fig. 7A, Table S7**). Only four genes (*cycA, relA, argW*, and *rnlB)* are categorized as primary important exclusively in this medium. Surprisingly, comparison with a list of known transporters in *E. coli*^38^ revealed that no transporter other than *cycA* is of primary importance in EZR, while 13 are of secondary importance. One possible explanation for this observation is the functional redundancy of the transporters. For example, both *uraA* or *rutG*^39^ genes are predicted to be responsible for uracil uptake but have a neutral effect on fitness in our screening. Meanwhile, the *upp* gene, which is required for uracil import by either one of those genes is important according to *i*Score.

## Discussion

The application of high-density transposon mutagenesis across 3 different growth conditions revealed a group of 412 common core important genes, complemented by groups of medium-specific important genes. The size and composition of this core group of important genes are coherent with other essentiality studies in *E. coli*^12,14^ and encode the main biological functions expected to be critical for bacterial life. On the other hand, we anticipated that the number of medium-specific genes would negatively correlate with the complexity of the medium. However, a total of 252 non-core primary important genes were identified in LB, of which 160 are unique to this medium, significantly more than the 116 and 4 genes exclusive to MOPS and EZR, respectively. This is striking as LB is based on yeast extract, i.e., whole *Saccharomyces cerevisiae* lysate, and is assumed to be the most complete of the three tested media^40^. However, optimal metabolization of LB is very complex, with cells sequentially depleting the most efficient amino acid available to harvest carbon^41^. We hypothesize that this process would require many more genes for a cell to be competitive in a growing population compared to one with a fixed and abundant carbon source, such as glucose. This supports the fact that the metabolism required for optimal proliferation in LB medium is highly intricate and fluctuates during growth, suggesting it may be ill-suited as a go-to medium for in-depth characterization, a point raised in several former studies^25,41,42^.

Overall, our results are consistent with those of other gene essentiality datasets, including other TIS-based studies^14^ and metabolic models^31^, while also providing additional refinement by identifying fitness-affecting genes through temporal variation analysis. We observed a few cases where genes determined to be essential by single gene knockout^12^ are not considered important in our assay. For example, *ubiJ, alsK, bcsB*, and *tnaB* have also been reported as non-essential by similar transposon-based work^14^. In general, discrepancies between our dataset and this latter work by Goodall *et al*.^14^ tend to involve smaller genes (Wilcoxon rank-sum test: Z-score = 8.76, p-value = 7.52e-22), with a median size of 489 bp compared to 810 bp for the entire *E. coli* gene set. Smaller genes typically exhibit fewer insertions and are more likely to fall just below or above the threshold of significance by chance. Another explanation for the observed differences between our competition-based approach and other single-gene inactivation studies is that some mutants may release growth-limiting nutrients into the medium. Strong competitive selection should eliminate most mutants with limited access to essential molecules^43^. However, nutrients required only in trace amounts, such as vitamins, might still be accessible in sufficient quantity to sustain mutants that would otherwise be unable to grow independently.

Genes identified as primary important are readily confirmed as having a major impact on fitness in predicted medium using competition assays (**Fig. S8**). For example, a Δ*tolQ* mutant, a core important gene, displays a substantial loss of fitness in both defined media. In contrast, deletion mutants of genes predicted to be medium-specific, such as *pheA* and *upp*, exhibit loss of fitness only in the predicted medium. Unexpectedly, some genes classified as neutral, such as *pdeB*, also exhibit a very slight fitness reduction, approximately one less division over ∼15 generations, compared to the parental strain. We would have expected this 1,551 bp gene to be classified as precarious or secondarily important, as this discrepancy cannot be attributed to its small size causing a false negative. This implies that other, yet undefined, phenomena may be at play during the selection process for certain genes.

An initial hypothesis concerning important gene regions was that occasional localized gaps of insertions could be caused by nucleoid-associated proteins (NAP), as suggested by a recent study^44^. This phenomenon could have been responsible for *MISS* regions. However, the previously reported insertion-depleted areas typically associated with NAPs were not detected in our dataset, despite the presence of the associated genes (*hns, mukB, tus*, etc.). Insertion sites showing an abnormally high read count, another type of potential artifact, tend to be common across various samples in all three media, suggesting an advantageous insertion site during the competitive enrichment and/or a Tn5 transposase insertion bias, as previously reported^45^. While the variety of sites that can receive transposons using Tn5 transposase is generally not limiting in TIS experiments, with nearly an insertion site every five base pairs in this study, exploring the precise in vivo insertion bias would be interesting. This includes factors such as NAPs, sequence specificity, and other more subtle causes, which could help improve the method by using this profile as a baseline to compare the post-selection insertion profiles.

Investigation of metabolic maps, such as the one for amino acid metabolism (**Fig. S7**) or the *uraA*/*rutG* redundancy, underscores that certain reactions can be catalyzed by more than one gene. This redundancy makes the importance of these genes hard to detect using approaches based on single-gene inactivation, such as single knock-out or HTTM. To develop a comprehensive knowledge base for thorough and predictive genomic simplification, a different approach involving whole-cell modeling^31,46^ or combinatorial gene inactivation^47^ will need to be devised to predict potential cases of synthetic lethality.

In addition to the 16 uncharacterized genes among the core important genes, there are an additional 146 genes with an uncharacterized status in the MOPS module and 60 in the EZR module (**Table S7**). Their varying importance across growth conditions could indicate a role in metabolism or its regulation. In addition, even well-characterized enzyme-encoding genes, although having an ostensibly clear function, may engage in moonlighting roles that are distinct from their annotated functions. This highlights the need for ongoing efforts to enhance gene characterization, even of a well-known organism like *E. coli*. Our medium-specific modules offer a practical design for precise, condition-specific engineering of this organism. Further investigations across a wider range of conditions, including different media and growth environments, would offer a more complete understanding of the specific contexts where each gene is important and help us learn more about their functions. This increased understanding will greatly enhance our ability to engineer microorganisms with specific capabilities for various biotechnological applications, while also expanding our fundamental knowledge about the function of numerous genes.

## Material and methods

### Growth conditions

Cells were grown in either LB broth (Miller), MOPS glucose or EZ-Rich (EZR) glucose. LB was prepared from commercial powder (BioBasic SD7003) at the following concentrations: tryptone 10 g/L, NaCl 10 g/L, yeast extract 5 g/L. MOPS and EZR were prepared following published recipes (https://www.genome.wisc.edu/resources/protocols/mopsminimal.htm and https://www.genome.wisc.edu/resources/protocols/ezmedium.htm). Exact composition of all media is available in **Table S3**. When required during the selection process, spectinomycin was added to a final concentration of 100 µg/mL.

### Transposon mutagenesis and library preparation

Transposon mutagenesis experiments and the preparation of sequencing libraries were performed as previously published^15^, with slight optimizations designed to increase efficiency. The main changes were 1) increasing the number of donors during the conjugation by ∼ 25 %; 2) changing the DNA polymerase used for the library amplification step for the SsoAdvanced Universal SYBR Green Supermix 2x (Bio-Rad); and 3) adding an additional depletion step during the library preparation to remove contaminating mutagenesis plasmid sequences from the genomic insertions. To do so, samples were treated with the AvrII restriction enzyme for 30 minutes at 37 °C after the ligation of the adapters to selectively digest the plasmid. Samples were then processed as previously described. The complete updated protocols are available on Protocol.io: https://www.protocols.io/view/httm-transposon-mutagenesis-dd3428qw, https://www.protocols.io/view/httm-gdna-extraction-dd3828rw, https://www.protocols.io/view/httm-illumina-library-preparation-dd4d28s6.

### Read processing

Paired-end sequencing reads were first assessed for quality using FastQC^48^ (v0.11.9) to evaluate per-base quality scores. Adapter sequences (ACTGTCTCTTATACACATCT) were then trimmed from both read pairs using Cutadapt^49^ (v4.9) and reads shorter than 10 nucleotides were discarded. Further quality filtering was performed using fastp^50^ (v0.23.2), applying right-end trimming with a sliding window of size 4 and a mean quality threshold of 10. Read alignment was performed onto the reference genome using bwa mem from BWA^51^ (v0.7.17) with a minimum seed length of 10. Output alignments were sorted and processed using samtools^52^ (v1.4), which includes the removal of low-quality alignments (mapQ ≤ 10).

Insertion site identification was performed by processing alignment files to produce per-position insertion counts, with single-read insertions discarded as noise. Feature assignment was achieved by intersecting insertion data with annotated genomic features using Bedtools^53^ (v2.30) while excluding features with low mappability, using previously computed with Genmap^54^ (v1.3). Gene-level insertion counts were computed by mapping insertion sites to genomic features. These profiles were then analysed using either Bio-TraDIS Toolkit^21^, which applies a statistical model based on prior distributions to classify genes as essential, ambiguous, or non-essential, or the sliding window analysis procedure described hereafter.

### Sliding window analysis procedure

To evaluate gene importance using the sliding window analysis procedure, each gene was scanned with overlapping sliding windows (bins) whose length was determined by the average distance between genomic insertions. Since most (> 80 %) of the genes are expected to be non-essential, we evaluated the number of bins containing significantly fewer reads than the global average. Assuming that insertion sites are random and that the distances between insertions follow a Poisson distribution, we found that in a bin seven times wider than the average insertion distance, there is approximately a 3% chance of having two or fewer insertions by chance (Fig. S9). Considering that not all insertions have the same number of reads associated with them, we used the number of reads associated with each insertion as a proxy for the quality of the insertion to discriminate between true insertions and background noise. For each bin, the total number of aligned reads was compared to the expected number of reads, corresponding to twice the median number of reads per insertion site in the sample. When above this threshold, the bin was considered *HIT* and thus probably not important. This means that a bin with a single insertion with more than twice the median reads count could make a bin *HIT* on its own, but it would only affect the gene’s score locally, while a bin with a few insertions having a single read each would be considered *MISS*. This approach provides resilience to both background noise and aberrantly high numbers of reads in single insertions that can randomly happen during this kind of experiment. A graphical representation of the process is available in **Figures 2 and S2**.

### Detection of important gene regions

Using the previously calculated bins, genes with a non-extreme *i*Scores (between 10 and 90) were scanned for stretches of quasi-consecutive *HIT* bins, meaning the longest stretch of *HIT* bin including at most a *MISS* bin. Genes exhibiting a streak of at least 20 % of their bins as *HIT* were selected for manual curation and inclusion as either a fully or partially important gene. The limit between the important region and the rest of the gene was determined as the bin having the highest ratio of *HIT* bins on one side and *MISS* bins on the other.

### Proteomaps

Proteomaps were generated using the Proteomap tool^33,55^ http://bionic-vis.biologie.uni-greifswald.de using a custom map template created from up-to-date *E. coli* KEGG Orthology data (https://www.genome.jp/brite/eco00001.keg). Unlike standard Proteomap templates where genes with multiple functions are assigned a function at random for plotting, genes with several functions are assigned suffixes (_1, _2, …) to allow mapping of all their known functions, and all known functions of a gene are displayed on the map using those aliases. The area assigned to each gene corresponds to the *i*Score squared, allowing for a greater separation from genes having a low number of insertions.

### Competition assays

Competition assays were performed to measure the relative fitness of a specific single-gene deletion mutant from the Keio collection^12^ against the BW25113 strain containing a kanR-GFP cassette inserted in *lacZ*^15^. For each tested medium, both the deletion mutant and the parental strain were precultured individually in the same medium, and an equivalent number of cells was mixed together based on their OD_600_ values. Mixed populations were then used to inoculate duplicate wells of a 96 deep well plate containing 1.5 mL of the tested medium, with an initial OD_600_ of ∼ 0.005. Deep well plates were then incubated at 37 °C and underwent a total of five daily passages under conditions similar to those of the HTTM experiment described at the beginning of the results section. To evaluate cell concentrations at the end of each passage, serial dilutions were performed in sterile 1X PBS, and the dilutions were fixed with a final concentration of 1% formaldehyde. Samples were then analysed using a BD Accuri C6 Plus flow cytometer (BD Biosciences) equipped with a 488 nm laser. The FSC-H channel threshold was set at 2500 to eliminate background noise, and the FL1-H (FITC) threshold was set at ∼ 200 to discriminate between the two populations. Fluidics were set to high speed, and a maximum of 100,000 events or 40 µL were collected for each sample. Cell populations were segregated based on FITC signal, distinguishing the BW25113 kanR-GFP strain from its competitor.

### Functional enrichment analysis

Statistical overrepresentation was calculated using the eco00001 Orthologs database from KEGG (https://www.genome.jp/brite/eco00001.keg) as a reference of functional terms. The 09150 Organismal Systems and 09160 Human Diseases categories were omitted as they are not relevant in *E. coli* K-12 strains. The enrichment analysis was performed using the hyperegom function from the SciPy^56^ Python library.

### *In silico* importance prediction using metabolic model

Flux balance analysis was performed on the iML1515 GEM using the COBRApy python module from the COBRA Toolbox^57^ with simulated MOPS or EZ-rich media. Genes whose inactivation lower the predicted fitness by more than 50% were considered important. Constraints used to simulate MOPS and EZ-rich media are the same as previously described^58^ and are based on the estimated medium composition available in **Table S3**.

### Time-resolved gene importance assays

Prior to gene read counts per base (read/bp) calculation, each sample’s read amount at every base position was capped at an upper threshold corresponding to the 95^th^ percentile, therefore negating strong swinging effects of single highly covered positions. Passages 1 and 2 were excluded since they harbored sparser transposon densities and higher artifactual plasmid reads. A pseudocount equal to the minimal non-null value of the dataset was added to the read counts per base for each preserved passage (3, 4, and 5) in both defined media (EZR and MOPS) for null values removal. Values were then normalized using geometric means for size factor calculation. These normalized values were transformed into logarithmic fold-changes (log_2_FC) using the P3 timepoint as the origin, in a similar fashion as Gallagher *et al*.^28^. The fold-changes were used to approximate depletion rate (log_2_FC/Passage) via standard linear regression.

To produce hierarchical clustering, normalized read counts per base were clipped at the upper extreme percentile. Along with the depletion rate, they were standardized (mean-centered and scaled to unit variance). As a driving factor, the depletion rate was upweighted 4 times compared to the three read counts per base. An embedding was produced using UMAP^59^ which was clustered using agglomerative clustering with SciPy’s^56^ default parameters and requesting 5 clusters. According to the depletion rate and final read count spectrum, clusters were attributed to a fitness label corresponding to their expected status (i.e., secondary important, precarious, and neutral). A visualization of the embedding with cluster labels is available in **Fig. S7**.

## Data availability

Raw reads of the HTTM experiments available as fastq files in the NCBI Sequence Read Archive under accession number: PRJNA1300301

A trackhub displaying all insertion sites at every timepoint for each medium is available at the following link : https://genome.ucsc.edu/cgi-bin/hgHubConnect?hgHub_do_redirect=on&hgHubConnect.remakeTrackHub=on&hgHub_do_firstDb=1&hubUrl=https://g-f2b62d.6d81c.5898.data.globus.org/Champie_2025/Champie_2025.hub.txt)

Supplementary Data are available online.

### Acknowledgements

We are grateful to the Centre de Calcul Scientifique of the Université de Sherbrooke for technical assistance. Access to computational resources was provided in part by Calcul Québec (http://www.calculquebec.ca) and Compute Canada.

## Author contributions statement

A.D.G. and A.C. performed all presented manipulations and sequencing.

A.C. and S.J., wrote the manuscript together with the support of A.D.G.

S.J. and A.C. performed the data analysis.

M.M.S. assisted with the data analysis.

D.M. assisted with data analysis and performed extensive curation of the text and figures.

A.C., S.R., A.D.G., S.J. and P.-E.J. designed the experiments and analysed the data.

P.-E.J., J.-P.C. and S.R. all contributed their expertise and resources to the advancement of the project.

