## Supplementary files for "Comparative Analysis of Gene Importance in *Escherichia coli* Across Growth Conditions"

**Supplementary Figures:**

- **Supplementary Figure S1: Number of genomic insertions per day in each tested medium.**
- **Supplementary Figure S2: Visual representation of the *i*Score analysis procedure.**
- **Supplementary Figure S3: Repartition of genes according to their *i*Score across all passages and growth media.**
- **Supplementary Figure S4: Determination of “essential” genes using exponential fitting of the non-essential gene population.**
- **Supplementary Figure S5: Validation of the *i*Score method and comparison with BioTradis.**
- **Supplementary Figure S6: Multidimensional classification of gene essentiality across passages based on read abundance and depletion dynamics.**
- **Supplementary Figure S7: Importance of the genes in the amino acid metabolism pathway in defined media.**
- **Supplementary Figure S8: Competition assay between mutants with different status and BW25113 wild type**.
- **Supplementary Figure S9. Probability of observing ≤2 insertions in bins of varying length.**

**Supplementary Tables:**

- **Table_S1_TnSeq_Statistics**
  - **Number of read and mapped insertion for each time point.**
- **Table_S2_Genes_iScore**
  - **iScore of all genes in all 3 media.**
- **Table_S3_Model_Predictions**
  - **Importance prediction made using he iML1515 metabolic model.**
- **Table_S4_Enriched_functions**
  - **Enriched KEGG functions in each primary important group of gene and in the modules.**
- **Table_S5_Partial_Genes**
  - **Description of genes harboring an important region.**
- **Table_S6_Data_Supp_slopes**
  - **Gene summary with read density, fold-change, and regression metrics.**
- **Table_S7_Modules**
  - **List of all genes composing both modules.**

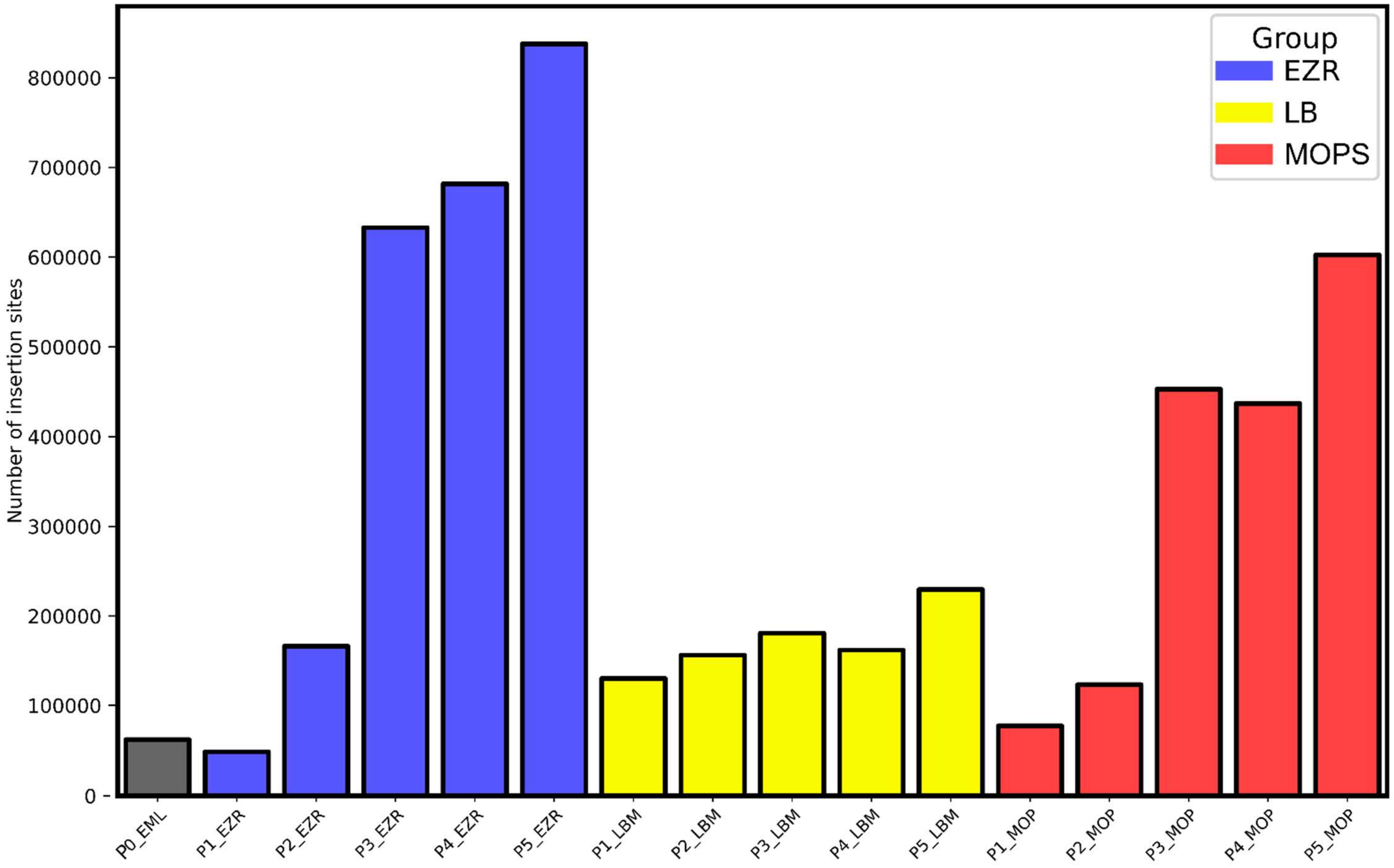

**Supplementary Figure S1: Total number of genomic insertions per day in each tested medium.** The Y-axis represents the number of total insertions in each sample after pooling all replicates on each passage. Passage "0" (P0) is pre-selection and is common for all media. EML: common to **E**Z-Rich, **M**OPS, and **L**B; EZR: EZ-Rich; LBM: LB medium; MOP: MOPS-glucose.

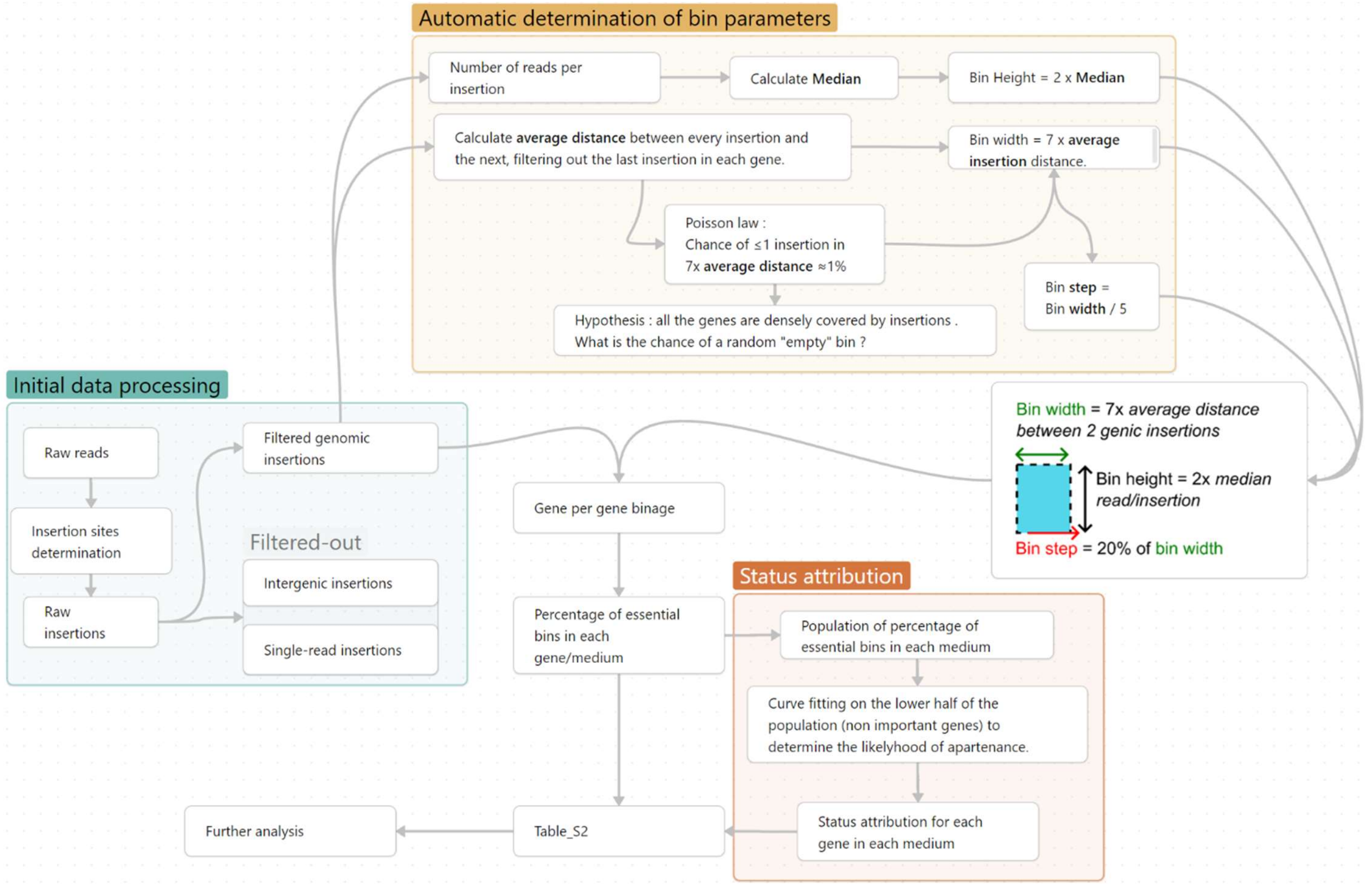

**Supplementary Figure S2: Visual representation of the *i*Score analysis procedure.** To evaluate gene importance, the procedure constists of three major steps: initial data processing, automatic determination of bin parameter, and status attribution. Raw reads used as input are pooled fastq files of all replicates for a given passage and experimental condition.

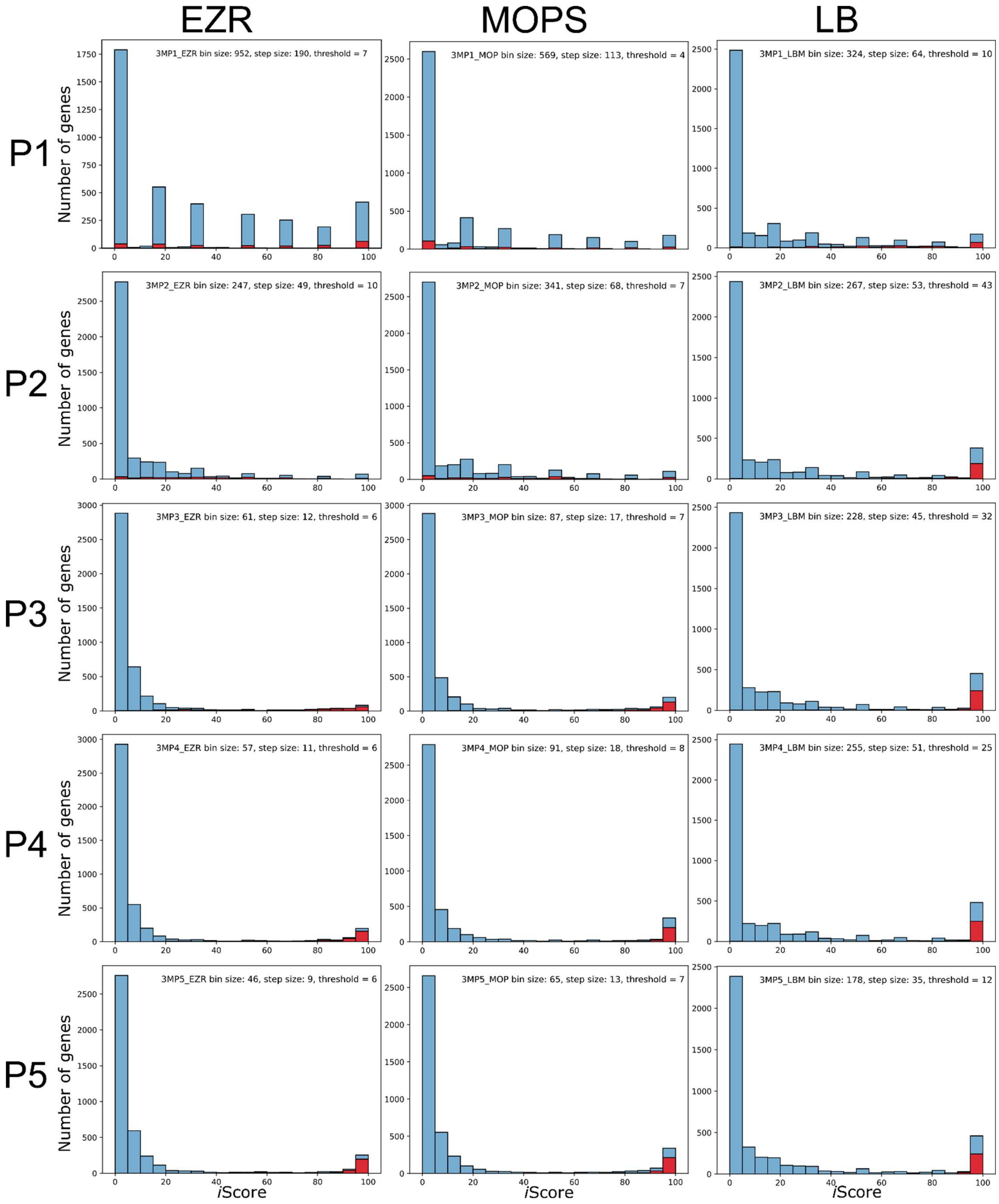

**Supplementary Figure S3: Repartition of genes according to their *i*Score across all passages and growth media.** Automatically determined bin size and number of reads thresholds are displayed in each plot. A subset of high-confidence important genes (genes determined to be important in all three media and with a size > 1000bp) are displayed as a red portion of bars for quality control.

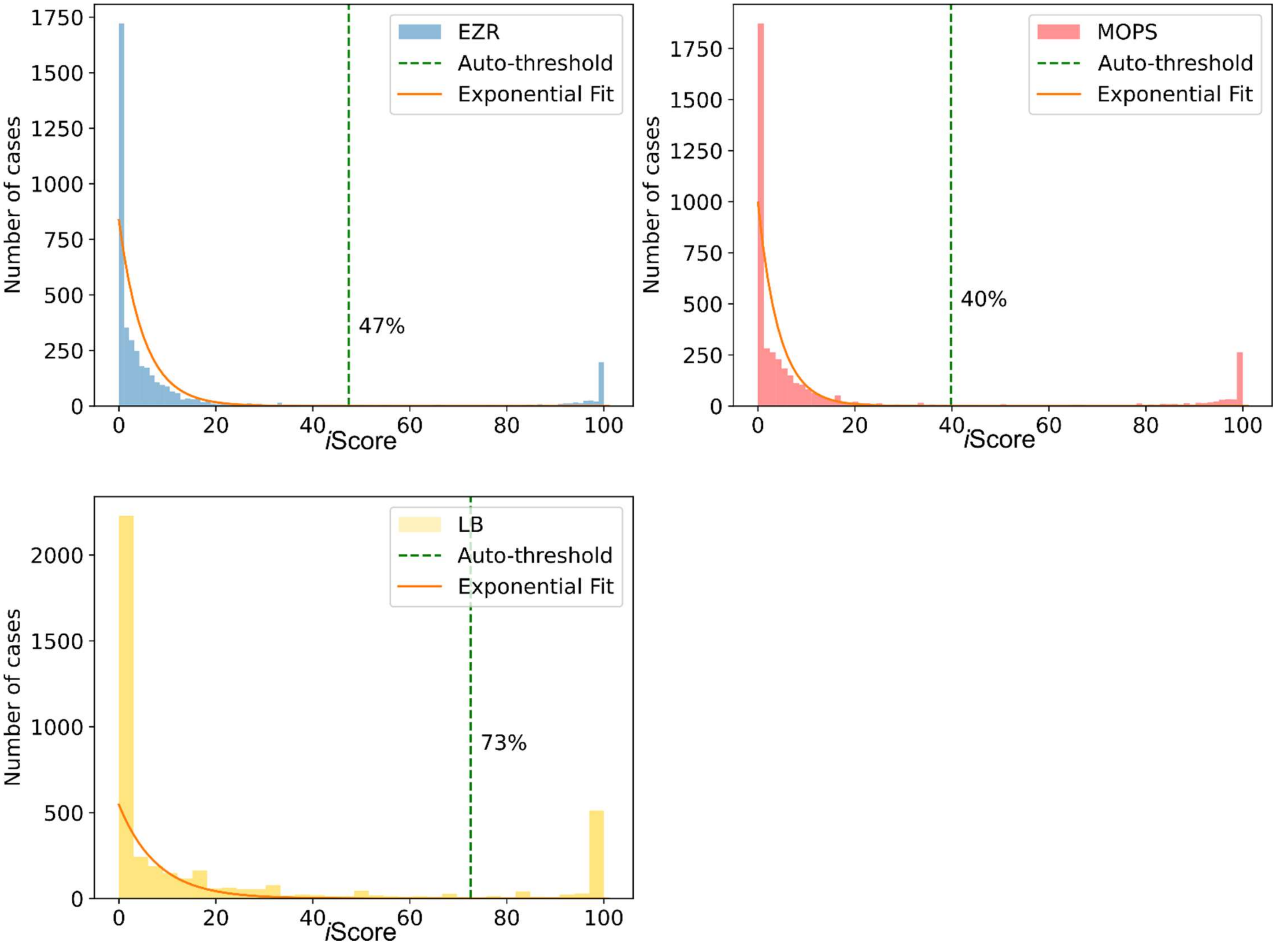

**Supplementary Figure S4: Determination of “essential” genes using exponential fitting of the non-essential gene population.** For each sample analyzed at passage 5 (from **Fig. S3**), the distribution of *i*Scores per gene exhibited a bimodal pattern. Using the left-hand population (iScore < 80), which represents genes of lesser importance, we estimated their distribution with a fitted exponentially decreasing curve. Utilizing this curve, we determined the percentage of bins required to have a <0.01% probability of association with the non-essential population, indicated by the green vertical bar. Genes with *i*Scores above this threshold (see caption for each medium) were labeled as “essential” genes. The relatively higher iScore required for classification as “essential” in the LB condition is caused by a lower insertion density in this condition which in turn causes the bins to be wider than in the other experiments and more prone to insertion noise.

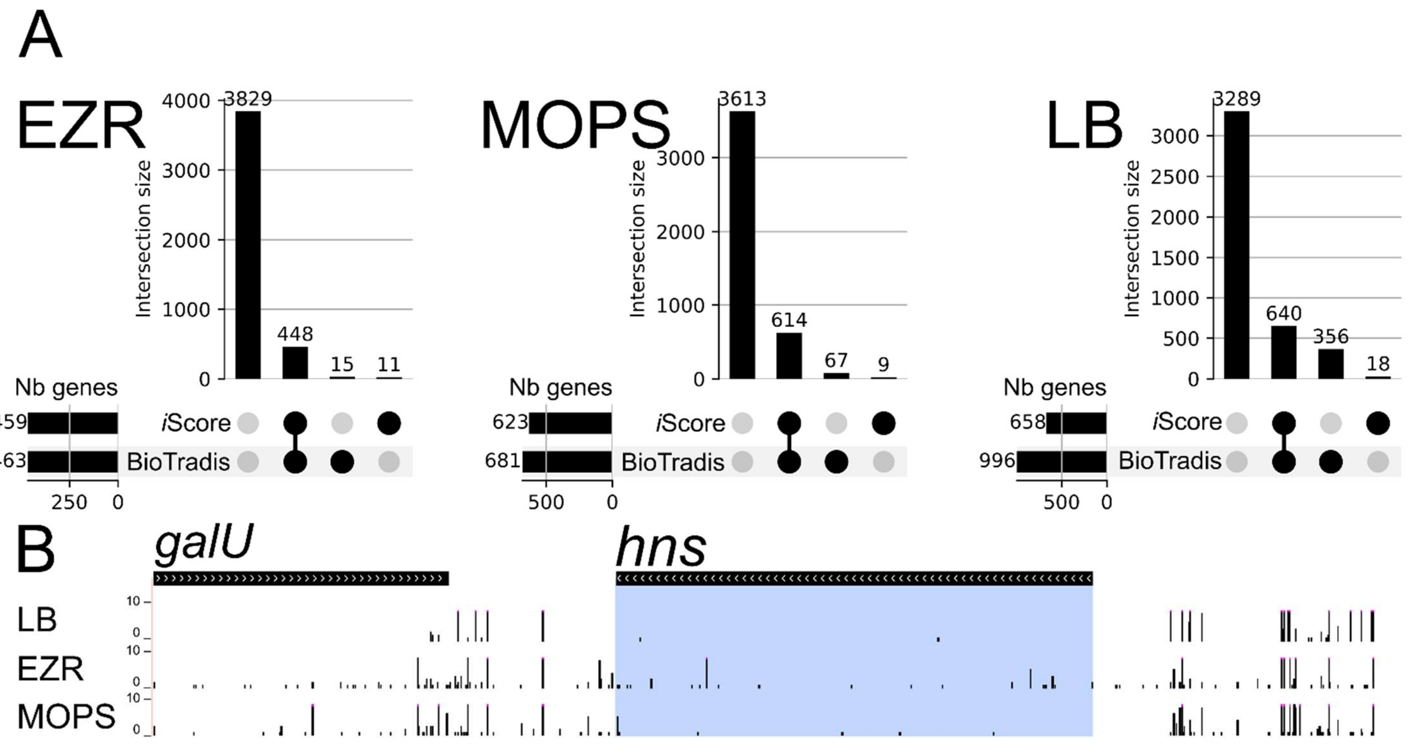

| C | Common | Considered Essential or Ambiguous by BioTradis Only | Considered important by *i*Score Only |
|---|---|---|---|
| **EZR** | **4261 (99.39%)** | **15**<br>*ybeD, sucB, rpmF, sdsN, dcd, ypeD, ibsC, yqgD, yrbN, trkA, ytiE* | **11**<br>*aceF, hemH, pflA, mukE, hns, sapA, ortT, ihfA, lapC, acpS, rnlB, higA, ilvX, efp, leuX,* |
| **MOPS** | **4259 (98.23%)** | **67**<br>*dnaJ, leuL, yaeP, ispA, ybaM, lipA, ybeD, rsfS, ybeY, ybgU, yljB, clpA, mukB, mepK, rmf, lpxL, rpmF, bhsA, mfd, ymfI, IS1414, rssB, pspA, hrpA, ortT, rnt, yoaL, zwf, dsrA, yeeA, asmA, cysP, ndk, iscU, acpS, rseA, rseD, ssrA, rnlB, micA, relA, rppH, ygfZ, yhaL, yrbN, rapZ, arcB, zapG, argR, tusB, tusC, secB, ysdE, ysdD, wecE, wecF, esrE, tatA, trkH, cpxR, metJ, aceK-int, hflX, ytgB, leuX, ryjB, yjjY* | **9**<br>*yahV, chiX, ychT, ruvA, yojO, yqcG, ilvL, cyaA, ytgA* |
| **LB** | **4117 (91.31%)** | **356**<br>*thrA, nhaA, carA, carB, fixC, apaH, pdxA, mraZ, aceE, mrcB, yadS, glnD, gloB, rnhA, prpB, lacZ, lacI, frmR, yaiY, queA, pgpA, yajQ, yajR, cyoE, cyoB, hupB, ppiD, priC, ybaK, purE, ybdZ, fepG, fepB, ybdD, rnk, citX, lipB, dacA, ybeZ, fur, ybfE, seqA, pxpA, nei, sdhX, mngR, ybgC, cpoB, nadA, aroG, galM, modC, pgl, bioF, moaB, dinG, fiu, moeA, deoR, rimK, ybjN, artQ, ybjQ, amiD, clpA, dmsA, dmsB, pflA, pflB, elyC, mepK, pncB, ssuD, uup, pqiB, ycbZ, ompA, hyaC, hyaD, cbpM, rutC, rutB, rutA, putA, efeB, opgG, opgH, yceK, pyrC, flgM, flgJ, flgK, flgL, fabF, pabC, mltG, ycfH, ycfL, thiK, nagZ, mfd, potB, pepT, phoQ, minE, dadA, dadX, dhaL, dhaK, ychF, dauA, narH, narI, rssB, oppC, oppD, oppF, yciU, yciC, trpD, rluB, sohB, pyrF, ycjT, tyrR, fnr, ldhA, paaH, paaI, paaJ, aldA, trg, opgD, ydcO, ydcV, patD, ddpX, hipA, lsrK, lsrA, lsrD, tam, rspR, ynfE, ynfG, dmsD, bidA, mlc, pntA, rstB, tus, fumC, manA, rsxG, dtpA, gstA, slyA, ydhI, ydhJ, sodC, ydhL, lhr, purR, punR, punC, mdtK, ydhT, ydhX, sufS, sufA, ydiJ, ydiS, aroH, btuD, ydiZ, astB, astD, astA, ynjA, ynjB, nudG, topB, selD, ydjA, yeaC, yoaE, yoaL, manX,* | **18**<br>*eyeA, ffs, appX, ymdF, ymgK, ychT, fnrS, rydC, sokB, yobI, rseX, yoeJ, yfiS, sibC, yhgO, yriB, ytgB, thrL* |

| | | |
|---|---|---|
| | | *mntP, mgrB, htpX, rsmF, yebW, ryeA, yebY, yebG, eda, zwf, pykA, lpxM, znuC, yebC, nudB, cmoA, cmoB, flhB, flhC, yedK, fliJ, fliN, yodD, dsrA, vsr, dcm, yedJ, tsuB, hisL, hisC, hisH, mdtA, mdtB, baeR, yegW, thiM, rcnR, mrp, mlrA, yehW, yohF, yeiW, bcr, rsuA, ccmG, ccmD, ccmA, napD, ada, ftp, yfaA, yfaE, glpT, arnE, menE, menH, ackA, pta, yfcE, cvpA, truA, pdxB, yfcJ, aroC, argW, fryA, cysZ, cysK, cysM, cysA, cysW, cysU, murQ, eutA, eutN, hda, purN, yfgM, hcaF, glyA, glnB, qseG, glmY, srmB, yfiM, kgtP, rluD, rnlB, nrdI, luxS, srlR, gutQ, hypC, yqcG, ygdG, ygdB, ppdA, ygdT, ygfX, gcvH, ygfB, tktA, gshB, yggS, hybD, exbD, ttdB, nfeR, ebgR, exuR, yqjD, yqjE, yhbQ, nlpI, rbfA, yhbE, rapZ, arcB, nanE, zapG, argR, rsmB, tusB, rpe, damX, gntR, ftsX, ftsE, rsmD, tusA, gor, baxL, mtlD, waaQ, rpmG, pyrE, spoT, ysdD, mnmE, mioC, rep, wecB, rffG, rffH, rffC, rffM, cyaA, yigA, mobA, polA, glnA, pfkA, hslU, thiS, purD, yjaA, pmrR, aspA, epmB, frdD, hflX, hflK, rnr, ulaR, pepA, ytiE, yjjV, serB* |

**Supplementary Figure S5: Validation of the *i*Score method and comparison with BioTradis. A)** Comparison of essentiality status attribution (black circles) after passage 5 data processed using either the BioTradis[1] methodology and the newly developed analysis procedure (***i*Score**). **B)** Example of a gene (*hns*) having discordant status between the two analyses in EZ-Rich (BioTradis = non-essential; *i*Score = important), where integration of the read/insertion counts likely leads to the differential status. **C)** List of all genes with a differential essentiality status between the two approaches.

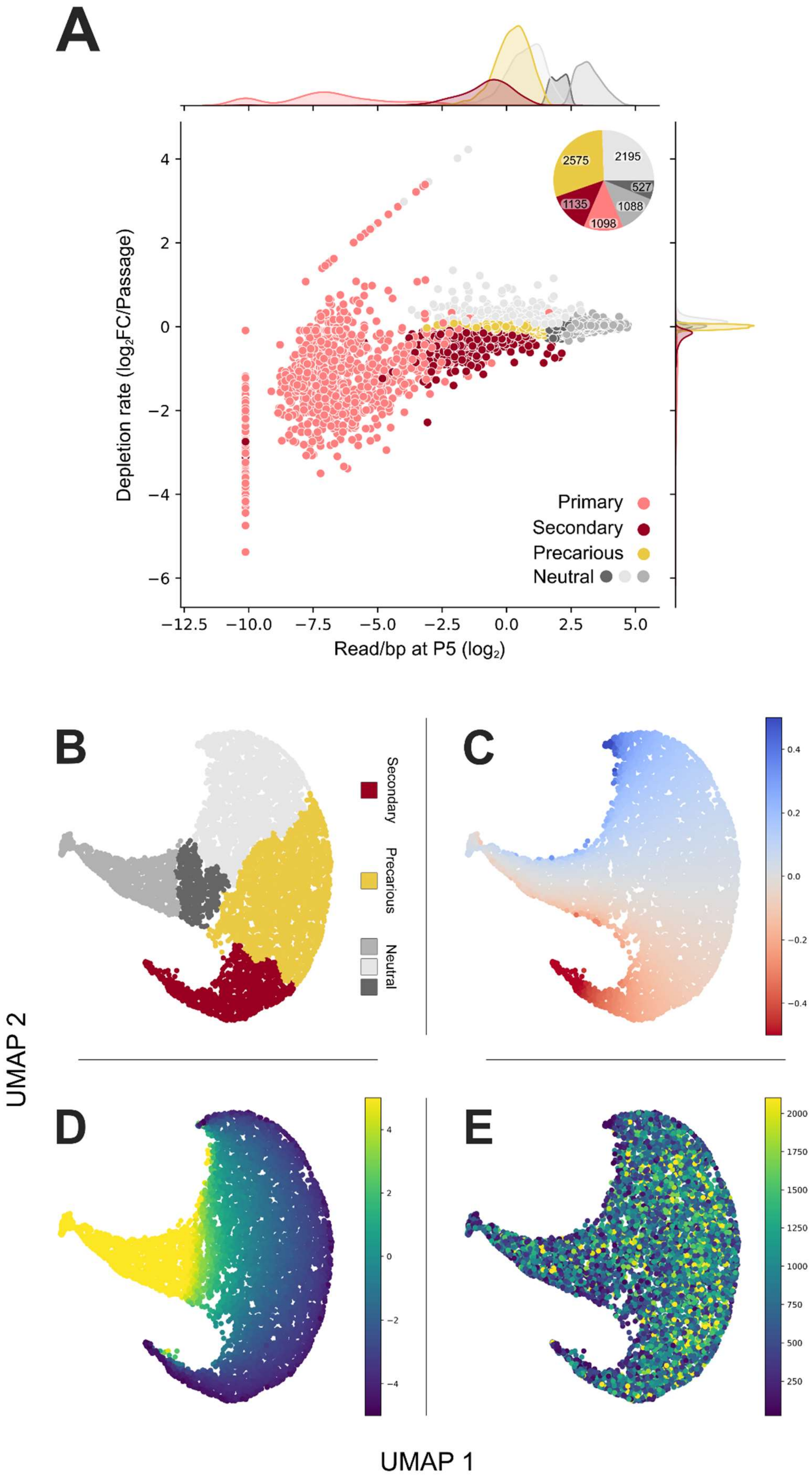
A
2575
2195
527
1135
1088
1098
Depletion rate (log2FC/Passage)
Read/bp at P5 (log2)
Primary
Secondary
Precarious
Neutral
B
C
D
E
Secondary
Precarious
Neutral
UMAP 2
UMAP 1

83

**Supplementary Figure S6. Multidimensional classification of gene essentiality across passages based on read abundance and depletion dynamics. A)** Depletion rate of all genes using normalized read counts across passages in both defined media vs their read count per base at P5. Values for both EZR and MOPS are plotted. Genes are colored according to hierarchical cluster assignments (primary, secondary, precarious or neutral). The upper right corner pie chart represents the total gene count in each cluster, both medium combined (see Fig. S6 for extended view including primary genes). Marginal density plots along the axes show the distribution of each metric across gene categories. The inset pie chart displays the absolute gene count per category. **B-E)** UMAP projections of genes based on their temporal profiles and abundance metrics, colored by **B)** essentiality class, **C)** $\log_2$ fold-change per passage (depletion rate) clipped at –0.5 to 0.5, **D)** scaled read count per base pair at P5 clipped at –5 to 5, **E)** gene length clipped ≤95% (suggesting gene length is not a primary driver of the observed clustering).

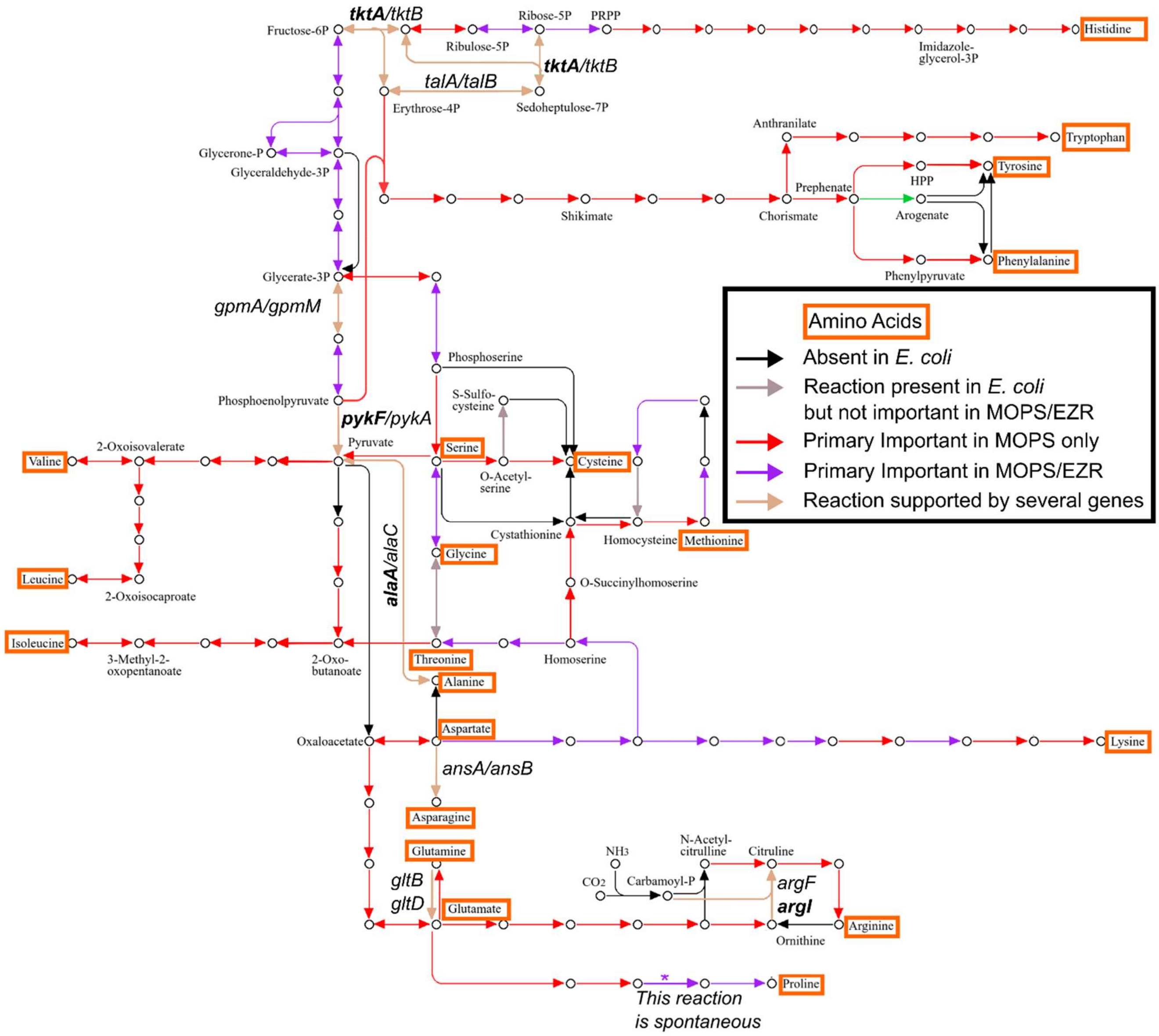

**Supplementary Figure S7: Importance of the genes in the amino acid metabolism pathway in defined media.** Based on the KEGG amino acid metabolism pathway map (eco01230), with manual recoloring. Reactions supported by multiple genes are annotated with the names of the known responsible genes. Among these, genes classified as secondary important in MOPS medium if any, are labeled in bold. Orange rectangles highlight amino acids within the pathway.

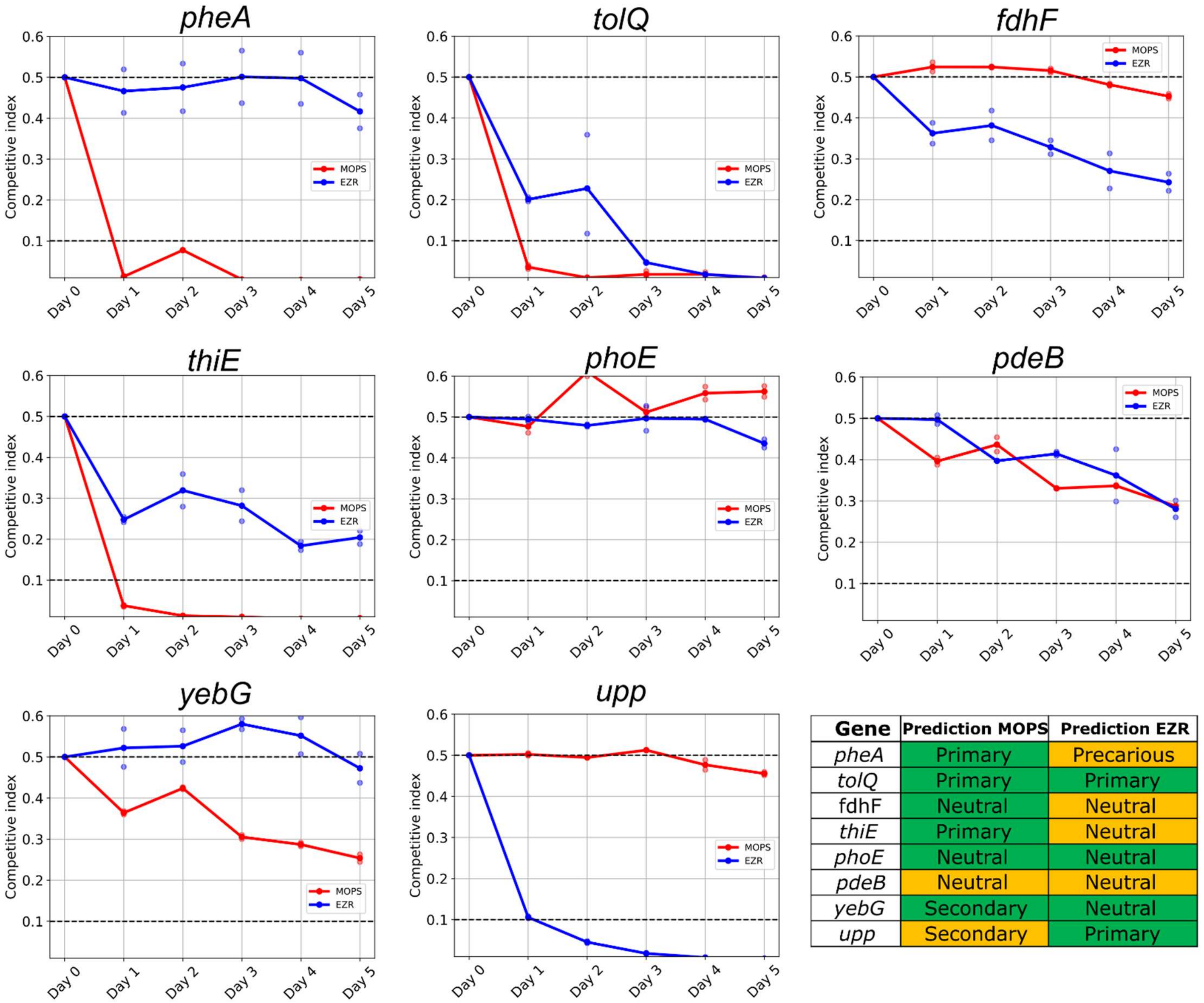

| Gene | Prediction MOPS | Prediction EZR |
|---|---|---|
| *pheA* | Primary | Precarious |
| *tolQ* | Primary | Primary |
| fdhF | Neutral | Neutral |
| *thiE* | Primary | Neutral |
| *phoE* | Neutral | Neutral |
| *pdeB* | Neutral | Neutral |
| *yebG* | Secondary | Neutral |
| *upp* | Secondary | Primary |

**Supplementary Figure S8. Competition assay between mutants with different status and BW25113 wild type**. After mixing equivalent amounts of cells using $OD_{600}$ (Day 0) cells concentrations were measured every day in duplicates using a FACS machine (Days 1-5). Y-axis represents the competitive index calculated as follows: (number of mutant / total population). Colors in the table represent the accuracy of the prediction, green = accurate, orange = mis-predicted.

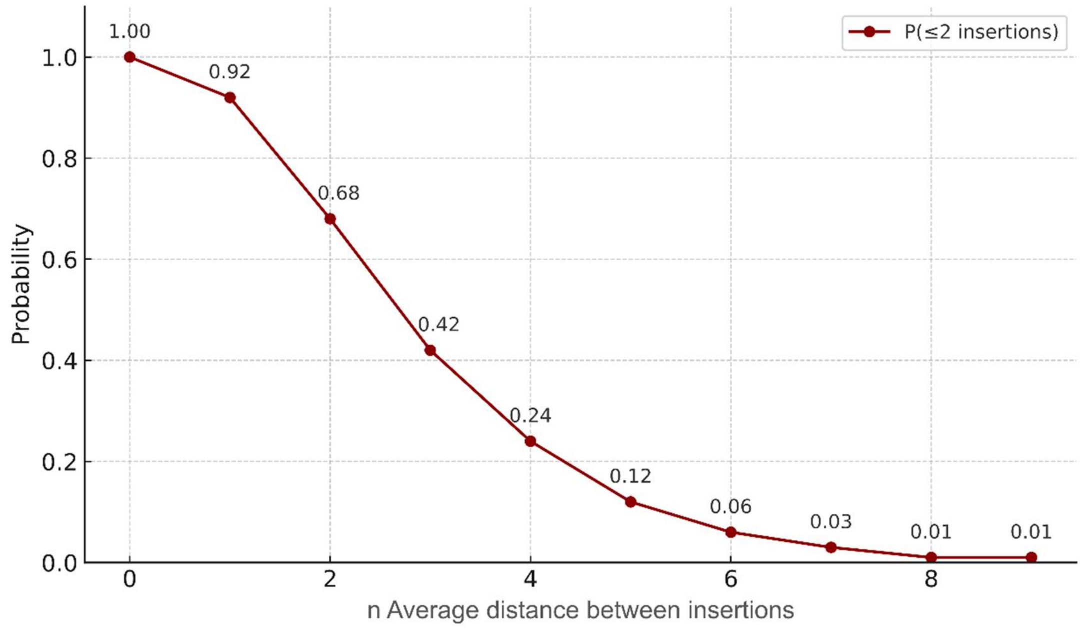

**Supplementary Figure S9. Probability of observing ≤2 insertions in bins of varying length.** The x-axis represents the bin size expressed as a multiple of the average insertion distance. The y-axis shows the probability of observing ≤2 insertions by chance in such a bin. Data point labels indicate the exact probability values.
